# Not all MDAs should be created equal – determinants of MDA impact and designing MDAs towards malaria elimination

**DOI:** 10.1101/793505

**Authors:** B. Gao, S. Saralamba, Y. Lubell, L. J. White, A. Dondorp, R. Aguas

**Affiliations:** Centre for Tropical Medicine and Global Health, Nuffield Department of Medicine, University of Oxford, Oxford, United Kingdom; Mahidol-Oxford Tropical Medicine Research Unit, Faculty of Tropical Medicine, Mahidol University, Bangkok, Thailand

## Abstract

Malaria remains at the forefront of scientific research and global political and funding agendas. Previous malaria models of mass-interventions have consistently oversimplified how mass interventions are implemented. We present an individual based, spatially explicit model of malaria transmission that includes all the programmatic implementation details of mass drug administration (MDA) campaigns. We uncover how the impact of MDA campaigns is determined by the interaction between implementation logistics, patterns of human mobility and how transmission risk is distributed over space. This translates into a higher likelihood of malaria elimination for areas with true prevalence under 3% with a faster implementation, in highly mobile populations. If populations are more static, deploying less interventions teams would be cost optimal and predicted to be as impactful. We conclude that mass drug interventions can be an invaluable tool towards malaria elimination in the right context, specifically when paired with effective vector control.

## Introduction

In Southeast Asia, and particularly the Greater Mekong Sub-region (GMS), *Plasmodium falciparum* transmission has decreased substantially over the last two decades (1, 2), setting the stage for pre-elimination scenarios, with all GMS countries committing to ambitious elimination timelines (3). Alignment of global funding bodies’ goodwill with sound national malaria control programmes is crucial for elimination timelines to be met (4, 5), but spreading artemisinin resistance creates a race against time before malaria becomes untreatable with currently available drugs (6, 7).

Vector control and early diagnosis followed by effective antimalarial treatment have been the mainstay of malaria control programmes, but modelling based projections indicate these approaches alone are unlikely to achieve falciparum malaria elimination before failing drug efficacy becomes an issue. Elimination will require more intensive measures to clear the infectious reservoir in asymptomatic populations, especially in the GMS where existing vector bionomics make vector control particularly challenging. The most abundant vector species in the GMS are exophilic (mainly bite outdoors), do not preferentially bite humans, and can bite quite early in the evening (8), rendering typical vector control measures such as insecticide treated nets (ITNs) and indoor residual spraying (IRS) sub-optimal.

Population wide interventions, including mass drug administration (MDA), are under consideration to clear the infectious reservoir in asymptomatic populations and potentially hasten progress toward elimination (9, 10). The proportion of the target population receiving these interventions (“coverage”) is believed to determine their success (4,11–13). This success can be considered at two spatial levels: global or local. Whilst malaria elimination campaigns have been carried out successfully in some countries or locally in specific regions (14–16), reintroductions of malaria from surrounding endemic areas are a constant threat (17, 18). The significance of mobile populations as a source of malaria transmission in the GMS has been emphasized in recent years (19–24). Prompt treatment of new clinical cases through malaria village worker (VMW) or village health worker (VHW) networks has proven to be an effective case management strategy (25, 26) and would be an essential barrier against malaria reintroduction.

We argue that the way in which mass interventions are deployed is what determines their success likelihood. The most efficient and effective roll-outs are laid on a solid community engagement foundation, thus ensuring subsequent adherence and coverage, while preventing malaria reintroduction from adjacent areas. Here, we model target areas as a collection of discrete villages (unit of intervention) and define coverage as the proportion of individuals receiving the intervention within a village and also as proportion of villages receiving the intervention within an area. Critically, we also simulate the minutia of mass intervention roll outs, with all its relevant deployment logistics thus assigning coverage a temporal dimension which measures the time it takes for all target villages to receive the intervention.

Conducting enough clinical trials to understand the interaction between all variables at play during a mass drug administration, as well as their individual and combined contribution to the expected outcome, is prohibitive. We developed a modular simulation platform that is customizable to any malaria transmission setting to provide realistic outcome predictions for local and global level interventions. The modules are the building blocks of an individual based, discrete time, spatially explicit, stochastic model, with explicit mosquito population dynamics and human population movements. We thus have villages with different mosquito densities connected by a human flow network, on which different interventions are deployed at different times (Figure 1). One particular innovation compared to previous published work (27–31) is the inclusion of very detailed logistical processes related to intervention deployment in the field. Whilst previously published models are extremely good at representing the biological processes underlying malaria transmission, some even making very realistic assumptions on how coverage increases over time (31), they fail to explicitly model how these interventions are carried out in the field. In practice, teams of workers visit villages sequentially one by one, usually spending 4-7 days to deploy one MDA round in each village, which is quite different from having an unlimited number of teams slowly treating everyone in the target population until a certain coverage is reached. Here, we investigate how the predicted impact of mass intervention strategies on malaria transmission changes when these logistical intricacies are taken into consideration. Our research questions are threefold: 1) what is the relevance of logistical implementation details to the outcome of mass interventions? 2) How fast does target coverage need to be reached for the strategy to be successful? 3) What are the key modulators of malaria elimination likelihood in a short time-frame? To offer strategic guidance to national malaria control programmes we also need to understand how the answers to these questions hinge on key features of malaria transmission in specific areas such as artemisinin resistance levels, population mobility networks, transmission heterogeneity over space, and seasonality patterns.

**Figure 1.**
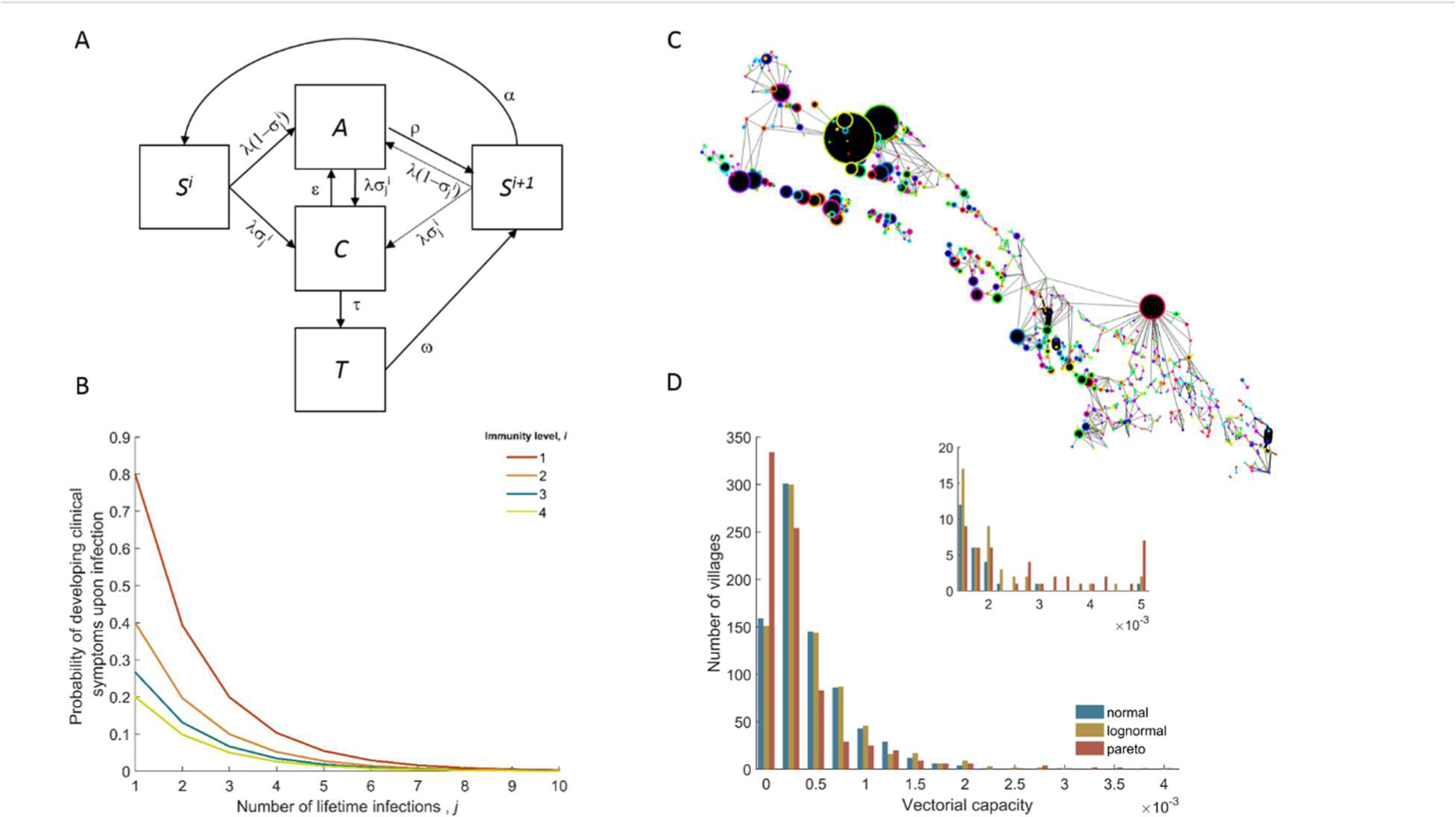
Model structure illustration. (A) Flow diagram representing the natural history of *Plasmodium falciparum* infections in human populations. Uninfected individuals (*S*) can be infected at rate *λ*, with the clinical outcome depending on the their immunity level *i*. Clinical infections (*C*) can be detected and subsequently treated at rate *τ*, or naturally subside into an asymptomatic parasite carrier stage (*A*). Upon recovery or treatment, individuals become susceptible with an added level of clinical immunity (*S*^*i*+1^). That clinical immunity can be lost at rate *α*. (B) Probability of developing clinical malaria depending on individual’s history of infection (cumulative number of infections) for each immunity level considered here. (C) Village connectivity network. Geo-located villages appear as circles the size of which is proportional to the number of people living there. Edge width reflects individuals’ probability of travel between connected villages. (D) Each village is assigned a specific mosquito density/vectorial capacity, with transmission heterogeneity over space characterized by three different distributions.

## Results

Initially, we simulated thousands of parameter sets that explore how a wide range of key transmission parameters (e.g. mean initial parasite prevalence, proportion of artemisinin resistant parasites) and logistical constraints – Table 1 – modulate the expected outcome of MDA campaigns. Figure 2 illustrates the sensitivity of the predicted proportional decrease in prevalence over 5 years to each parameter. Clearly, the number of MDA rounds and the initial mean prevalence across all villages are critical covariates when predicting MDA outcome. Also quite important seem to be the distributions characterizing how malaria risk is distributed over space. Artemisinin resistance spread is quite sensitive to the same covariates and additionally to the number of intervention teams and the intensity of human population mobility (Figure 2 figure supplements 4,5).

**Figure 2.**
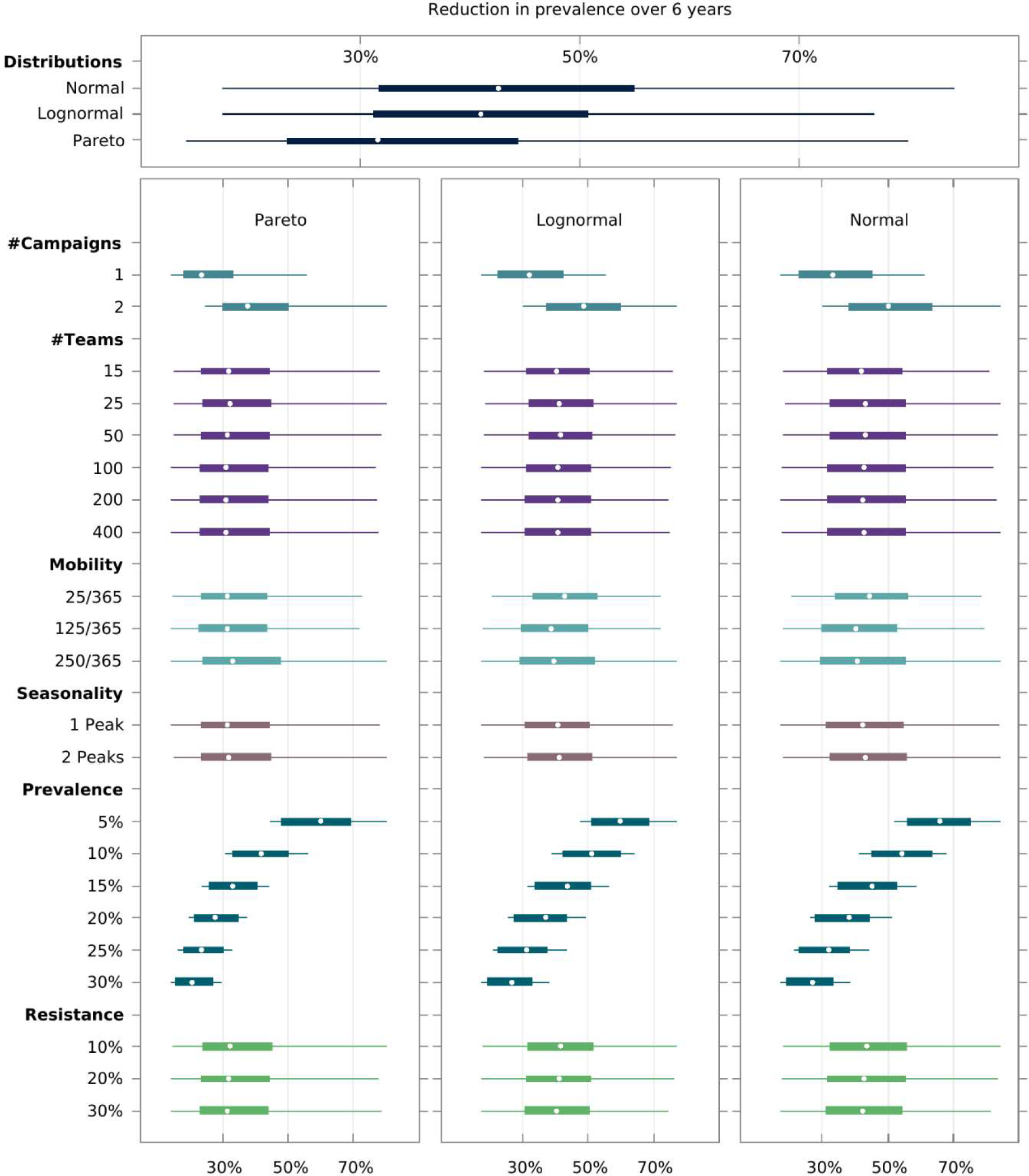
Multivariate sensitivity analysis of the predicted intervention impact on malaria prevalence and artemisinin resistance. The box plots show the median and interquartile ranges of the proportional reduction in malaria prevalence for all simulated parameter sets. Each parameter set consists of a combination of initial mean prevalence (*Prevalence*), initial proportion of artemisinin resistance parasites (*Resistance*), population mobility (*Mobility*), number of teams deployed in the field (*# Teams*), number of MDA campaigns (*# Campaigns*), and number of transmission peaks per year (*Seasonality*). An overall mean and interquartile range for the effect of transmission heterogeneity, independent of any other parameter, is displayed on the top panel. The reduction in prevalence is evaluated as the proportional difference in the integral in prevalence in the 5 years following MDA relative to the 5 years preceding MDA.

**Table 1.** Factors explored by the model and their respective sets of values.

| Factor | Meaning | Values |
| --- | --- | --- |
| Teams | Number of teams performing prevalence surveys and distributing ACTs simultaneously. Translated into coverage speed, *i.e.*, the time it takes to perform one MDA round in a region of 1,000 villages. | 15, 25, 50, 100, 200, 400 |
| Prevalence | Mean initial malaria u-PCR prevalence across all villages. | 0.5, 0.1, 0.15, 0.2, 0.25, 0.3 |
| Resistance | Mean initial proportion of parasites which are artemisinin resistant. | 0.1, 0.2, 0.3 |
| Distribution | Underlying spatial heterogeneity of malaria transmission, manifested by differential mosquito density distributions. | Gaussian, Log-normal, Pareto |
| Mobility | Describes the intensity of population movement in the general population. Indicates the per person average daily probability of staying overnight in places other than their home village. | 0.068, 0.342, 0.685 |
| Campaigns | The number of MDA Campaigns | 1, 2 |
| Seasons | The number of mosquito peak seasons | 1, 2 |

We found that there is an intricate relationship between the optimal timing of MDA campaign start, its implementation logistics, and malaria seasonality patterns. Deploying a higher number of MDA teams will yield a higher likelihood of reaching malaria elimination within 2 years, only when population mobility is high – Figure 3. Using a smaller number of intervention teams is predicted to be advantageous in more static population, especially when there is only one annual transmission peak and when the MDA start is delayed to day 60 (instead of the default start at the beginning of the calendar year). In settings with 2 malaria seasons per year, the first transmission peak occurs earlier in the year, making the faster 400 team implementation a better option in general. The only exceptions are very static populations in which two MDA campaigns are deployed. We should note that overall, a slower implementation is preferable, as there is little difference in the likelihood of elimination within 10 years (Figure 3 – figure supplement 1), especially for the single peak seasonal profiles. For the two peak scenarios, a higher number of teams would be beneficial as prevalence increases from 1%. When addressing how to maximize the chances of reaching elimination within a short time span by implementing an MDA strategy, we found that different transmission heterogeneity distributions incur quite different prospects – Figure 4. The likelihood of reaching elimination is strikingly different when comparing the Pareto distribution (most skewed) – Figure 4 supplement 2 – with the other two, except when an extremely efficient vector control program is carried out for a couple of years. Coupling vector control with an MDA campaign vastly improves the chances for elimination across the low prevalence settings explored here, and the longer vector control can be sustained the more likely elimination becomes. Once again, we retrieve a significant correlation between population mobility and number of MDA teams, with faster MDA implementations being preferred when the human population is more mobile.

**Figure 3.**
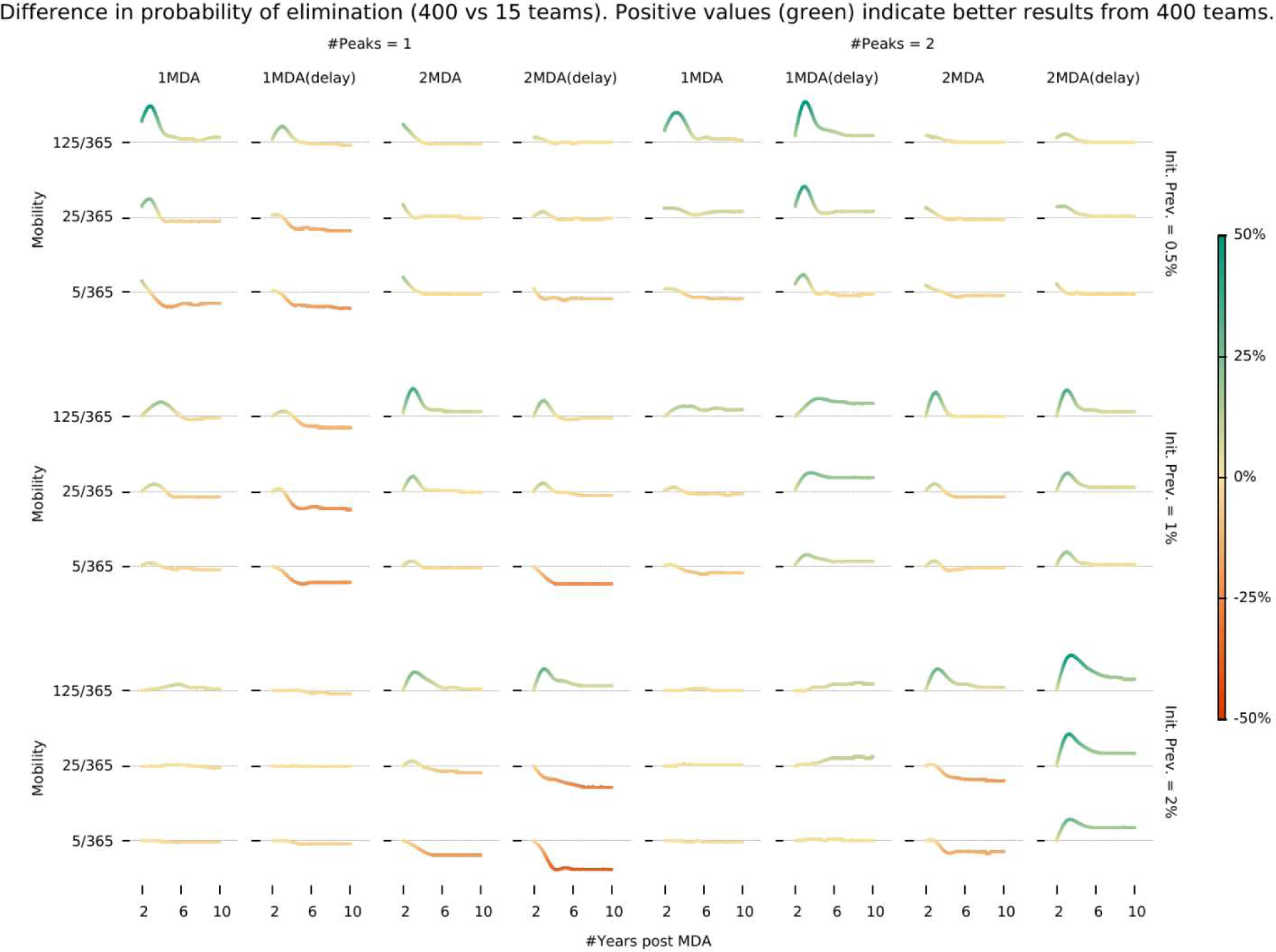
Intervention implementation speed in different epidemiological contexts. Demonstrates under what conditions using 400 implementation teams is preferable over 15 teams when deploying an MDA campaign. We investigate different epidemiological contexts, characterized by different prevalence and seasonality profiles can be accounted for in deciding the appropriate campaign start day (delay) when maximizing the chances for malaria elimination.

**Figure 4.**
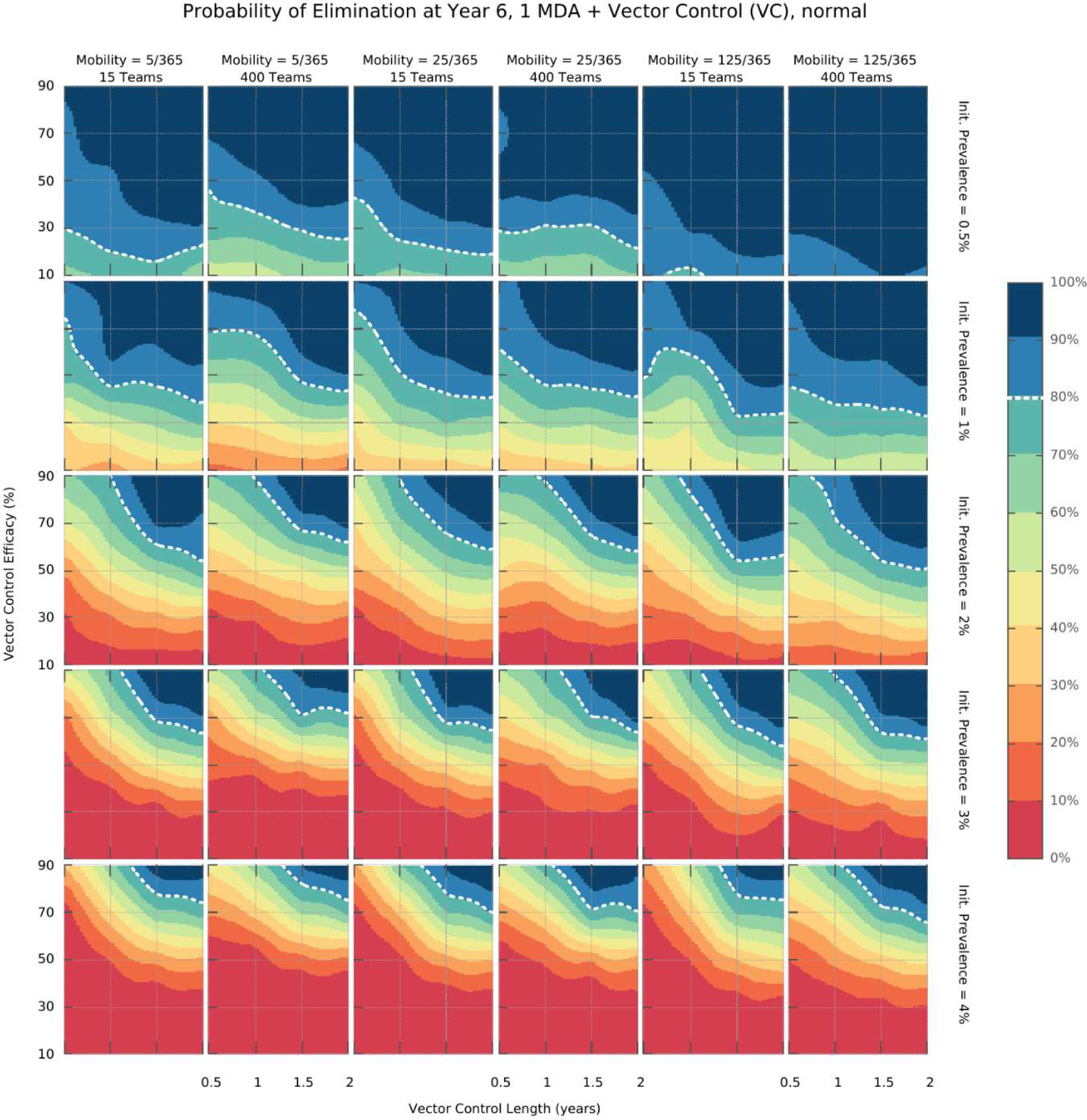
Elimination probability surfaces. These surface plots show the proportion of simulations (out of 100) in which elimination was achieved within a 5-year time horizon. Hypothetical interventions that decrease the vectorial capacity by a proportion given in the y-axis are maintained for a period of time defined in the x-axis. These transmission blocking interventions are layered on top of a global MDA strategy consisting of 3 ACT rounds in all villages and including widespread village malaria workers. Different panels give different combinations of mean initial malaria prevalence, human population mobility and MDA implementation speed. The white dashed line represents the 80% likelihood of elimination contour line.

Finally, we explored the value of targeting the top 10 or 20% of villages (sorted by vectorial capacity) and compared its predicted outcome with a full MDA campaign – Figure 5. We confirm that the log-normal and gaussian distributions explored here produce the same elimination likelihood profiles. For very efficacious vector control strategies, the vectorial capacity in the transmission foci will be greatly reduced, causing a greater drop in mean vectorial capacity across all villages in the more skewed distribution (where most villages have negligible numbers or no mosquitoes), compared to the others. This causes the likelihood of elimination in settings with a Pareto distributed risk of infection to be greater on the long-term under those circumstances. Once again, sustaining the vector control for longer, greatly improves the expected outcome (Figure 5 – figure supplement 1.

**Figure 5.**
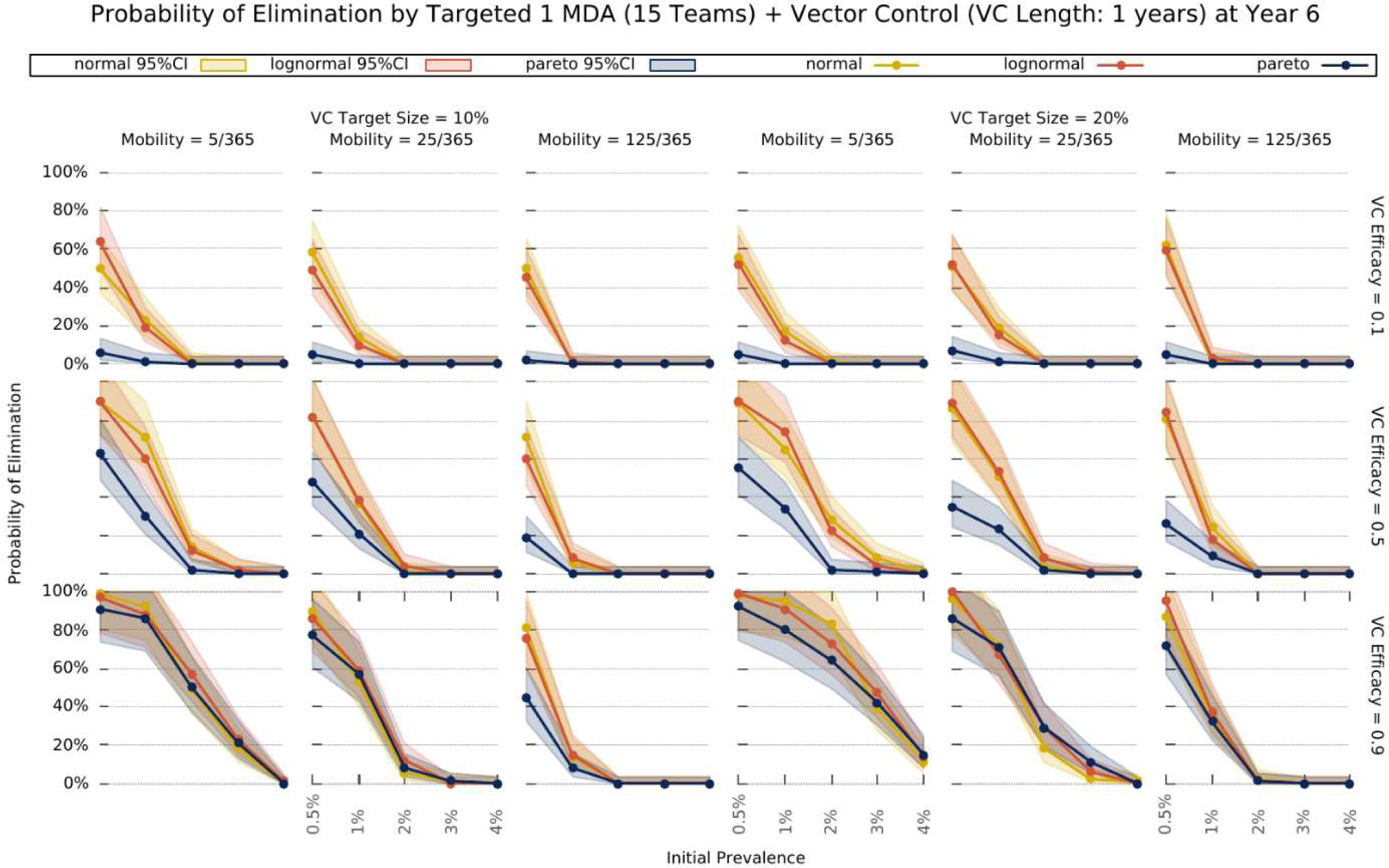
Elimination probability with a targeted approach. Illustrates the likelihood of elimination within 5 years of an elimination strategy consisting of 3 MDA rounds and a vector control strategy sustained over 1 year. We compare elimination prospects across different prevalence levels, human population mobility, and transmission heterogeneity over space. Vector control (VC) efficacy refers to the coefficient by which vectorial capacity is reduced for the duration of the intervention. The vector control target sizes refer to the quantile of villages, sorted by descending vectorial capacity, target with the intervention.

## Discussion

A series of key interacting features of the transmission-intervention system emerge when intricate logistics are incorporated in spatial-temporal transmission dynamics. Mapping MDA campaign expected outcomes to a specific malaria endemic setting is a complex multivariate problem. Here, we elucidate the way in which the most critical interactions determine MDA success:

### Operational strategy design

Mass intervention strategies rely on a detailed protocol defining the proportion of villages targeted, the target population in each village reached (usually termed target coverage), and the number of intervention teams deployed (determining the speed with which all villages are covered). Unsurprisingly, the chosen number of MDA rounds is the most significant intervention outcome determinant (Figure 2). This is intuitive in a scenario where treatment failure due to drug resistance is not a serious issue. Assuming 80% of the target population receives each MDA round, and independent coverage between rounds (meaning the likelihood that someone adheres to round 3 e.g. is independent of their uptake in rounds 1 and 2), by increasing the number of MDA rounds, we are decreasing the proportion of the population not treated with at least one round of ACT. Even if adherence and compliance are correlated, increasing the number of rounds would assure individuals that received treatment would be less likely to become infectious, or be infectious for long, if infected via untreated individuals. Indeed, additional MDA rounds provide a powerful tool to disrupt any resurgence in transmission following the typical 3 round MDA campaigns. The likelihood of elimination being achieved is substantially higher for 2 sets of 3 MDA rounds compared to 1 (Figure 3). Interestingly, transmission heterogeneity, described by different mosquito number distributions over space, and initial mean parasite prevalence in the human population show up as distant second and third in terms of model sensitivity (Figure 2 – figure supplements 1 and 2). Of note, MDA strategies on their own are not predicted to achieve elimination unless mean malaria prevalence is at very low levels (under 5%) – Figures 2 and 3. Coverage speed, increased with a higher number of intervention teams, is programmatically beneficial only when elimination is achievable and in scenarios where MDA is deployed in well-connected populations – Figure 3. Counterintuitively, in all other scenarios, a slower MDA implementation is optimal (Figure 2 – figure supplement 3, Figure 3 – figure supplement 1). When the number of MDA teams is highest, all villages receive the first round within a couple of weeks, meaning a very large proportion of the population will be under treatment simultaneously. Whilst that translates into the largest possible increase in the likelihood of elimination, any resistant infections will have a large selective advantage at that point, causing resistance to spread (Figure 2 – figure supplement 5). Interestingly, this effect is less pronounced in populations with high mobility, due to a dilution effect described below. A slower deployment of MDA campaigns is then generally preferable in low mobility populations (here defined as settings where individuals spend on average less than 25 nights per year somewhere other than their home) due to a slower buildup of resistant infections (Figure 4).

### Transmission topology

Defined by the magnitude of transmission heterogeneity over space combined with the level of mixture between sub-populations through human movement. As mentioned above, spatial transmission heterogeneity has a dramatic effect on predicted outcomes, with more skewed distributions (suggesting most malaria infections occur in a few villages) presenting a challenge for control (Figure 4 – figure supplement 2). We investigated a spectrum of population mobility patterns which at one extreme consists of a set of isolated transmission foci (thus low population movement), where infections in each village are almost exclusively locally acquired (mosquitoes infecting someone will have had acquired that infection from someone else living in the same village). As population mobility increases, these foci become more and more connected, eventually merging with each other in the upper extreme of the spectrum, onto one large homogeneously mixed population (where mosquitoes infecting a person could have acquired that infection from anyone else).

In low connectivity populations, the likelihood of resurgence in villages where MDA achieved local elimination is low, since there are only very sporadic introductions of parasites, typically insufficient to reseed endogenous transmission. In these settings, implementation speed should be sacrificed, and a low number of intervention teams deployed, to minimize resistance spread. In populations with high population movement, speed of implementation becomes more important (Figure 3) due to the propensity for recently eliminated intervention units to be reseeded by its neighbors, leading to local resurgences.

### Seasonality

The theoretically best timing of MDA campaigns relative to the malaria season peak have been investigated in (32). Critically, those models do not incorporate implementation logistics in a detailed manner, either simulating instantaneous MDA deployment in all villages, or having a synchronously increasing global coverage until a target coverage is reached. Here, we have explicit intervention teams that deploy sequential MDAs, one village at a time. This is much closer to reality in the field and creates an added dimension when comparing MDA start with malaria seasonal peaks. A very slow implementation that starts 3 months prior to the peak might only have reached a 50% coverage by the time transmission intensity hits the peak, whereas a very fast MDA deployment might start 1 month prior to the peak and end before it.

When there is only one transmission peak during the year, a slower MDA implementation seems to be preferable (Figure 3), especially if the start of the MDA is set to start one month prior to the peak in vectorial capacity (not to be mistaken with the season malaria incidence peak which occurs later) instead of starting at the beginning of the year. It seems delaying the start of MDA campaigns improves the likelihood of malaria elimination compared to a start at the beginning of the year when a lower number of MDA teams is used (Figure 3 - supplement 1). This is mostly due to how sensitive near instantaneous MDA campaigns are to the timing of the seasonal peak. In a single annual peak setting, where the incidence peak is at day 140, a near instantaneous MDA would end (all 3 rounds) 70 (for no delay) or 10 (with delay) days prior to the peak. Given the general cosine function simulated here, it seems a longer implementation lasting the whole duration of the high transmission season is optimal. For the two annual peaks scenario, a fast implementation of MDA (covering the whole of the first annual peak) seems preferable.

### Dilution

We uncovered an interesting trade-off between population mobility, transmission heterogeneity and number of MDA teams that results in unexpectedly high predictions for intervention impact in highly mobile populations. This is due to a diluting effect, rooted in the sharing of parasite pools between high transmission foci and very low transmission villages, which is particularly relevant in the post MDA rebound period. Granted high population mobility, after the parasite pool is greatly reduced through MDA, the few infectious mosquitoes in transmission foci are likely to bite migrants, which upon return to low transmission villages, are unlikely to transmit those infections onwards. Increased mobility decreases the proportion of endogenous infections in each village, which consequently increases the radial impact of local interventions, thus also generally contributing to the dilution of artemisinin resistant emerging infections (Figure 2 – supplement 5). In a scenario where village A with an extremely low vectorial capacity is very close to a high transmission village B and there is intense population movement between villages, it is likely that infections in people living in village A are almost exclusively acquired when they visit village B. Thus, an MDA in village B will have a positive knock-on effect on the incidence of malaria in village A.

### Drug Resistance

We observe that increased transmission heterogeneity is detrimental to intervention success (Figure 4 – figure supplement 2), through a mechanism whereby high transmission foci provide a niche for resistance spread which is promoted by drug pressure incurred through multiple MDA rounds (Figure 4, Figure 2 – figure supplement 4). Interestingly, this effect is more pronounced for populations with the lowest mobility for all considered spatial heterogeneity distributions. This is due to the dilution effect mentioned above, causing artemisinin resistant parasites expanding in high transmission foci to be diluted across other villages, given a sufficiently high population movement. If the population is very static, however, resistant parasites can gradually outcompete sensitive ones in transmission foci receiving multiple MDA rounds. Drug resistance is also a key driver of why more heterogeneous topologies are predicted to have a lower MDA impact for low mobility populations (red lines in Figure 2 – figure supplement 5). Thus, any concerns regarding the enhancement of artemisinin resistance spread with MDA hinge on the transmission heterogeneity of the setting in which MDA is deployed. This entails serious consequences for the likelihood of malaria elimination with an MDA approach in populations where resistance is already established.

Although the elucidation of these intricate relationships is of great scientific interest, National Malaria Control Programmes (NMCPs) might find this exploration to be devoid of applicability, specifically those concerned with drug resistance issues. We have demonstrated that more MDA rounds translate into higher likelihoods of malaria elimination but also show how resistance is very likely to increase dramatically if elimination is not achieved. To explicitly inform policy decisions of NMCPs aiming to eliminate *P. falciparum* malaria in a short time frame, we provide insights into integrated strategies that combine vector control interventions with a minimal number of MDA rounds:

### Intervention layering and elimination prospects

While MDA strategies consisting of only 3 rounds of ACT are unlikely to interrupt transmission in all but very low prevalence (<2 % all-age true prevalence) settings, the village malaria worker (VMW) network providing operation support to MDA campaigns does provide a great foundation for additional interventions to be more easily deployed. Combining a complete VMW network with 3 rounds of MDA at 80% coverage and imperfect vector control strategies, we predict malaria elimination can be reached, provided the initial parasite prevalence is sufficiently low (under ∼2%) – Figure 4. More than a theoretical possibility, elimination, even when in the presence of artemisinin resistance, has been demonstrated to be possible (33). This bistability phenomenon, first proposed for malaria a decade ago (34) and since revisited (35), provides the theoretical foundation for the determination of the minimum intervention effort sustained over a defined period of time, after which all intervention measures can be relaxed and elimination is still reached (provided clinical case management remains effective). We thus explored the prospects for elimination with realistic intervention packages coupling vector control with MDAs using our simulation platform – Figure 5. We find that initial mean prevalence is a key determinant of success likelihood and determines the minimum effect size required from the vector control component, for elimination to be reached. We highlight how logistical constraints combine with human movement patterns to modulate the likelihood of an intervention strategy’s success. Clearly, when population mobility is high, elimination becomes likely even with low vector control effect sizes, if MDA is done near instantaneously (400 intervention teams). A much higher vector control effect size would be required if MDA implementation is significantly slower. Conversely, if the human population is static, a slower MDA implementation would increase the likelihood of elimination for the same vector control effect sizes. This illustrates well the value of information of missing dimension of simulation models that convert the highly complex logistics of a global MDA administration into an instantaneous process.

### Targeting interventions

Over the last few years, MDA interventions have moved towards a focal approach, where only a proportion of individuals within a village (36), or a proportion of villages in the target area will receive treatment (37), to ease drug resistance spread concerns, and to minimize the number of ACT doses given to uninfected individuals. The purpose is to implement MDAs in high transmission foci only, thus bringing mean prevalence across the whole population down very quickly, and then relying on good clinical case management and vector control interventions to eventually reach elimination. The logistical implementation is also simplified, and its associated costs minimized, since only a fraction of MDAs are performed. We explored the likelihood of reaching malaria elimination within 5 years by implementing a targeted elimination program combining 3 rounds of MDA with different vector control intensities and found that the duration and effect size of the vector control component greatly influences the prospects for elimination. Interestingly, increasing the target size (number of villages receiving the intervention), minimizes the differences across transmission heterogeneities.

In fact, the most skewed distribution offers better elimination prospects for higher prevalence settings where intense vector control (VC efficacy = 0.9) are put in place. The nature of the distribution provides the basis for this effect. Given that most villages have a negligible vectorial capacity, transmission is sustained by a few high transmission foci. If those are targeted efficiently, you can expect to disrupt transmission more than in settings where the distribution of mosquitoes over space is more homogeneous.

We have refrained from doing a cost-effectiveness analysis, since we do not have good information on most of the unit costs, which are currently being assessed in different settings, and are likely to be quite variable across countries in the GMS. Any recommendation and cost-effectiveness analysis would have to be tailored to each specific country/area. We also have not extensively addressed synchronous migration patterns but have included long term and seasonal migration events in the simulation platform. The sensitivity of the model’s predictions to these types of migration is much lower than to the general population migration explored to great lengths throughout. This is, in all likelihood, a result of the low proportion of seasonal or long-term migrants in the overall population. In settings where migrants constitute a more considerable (>20%) fraction of the population, the predicted impact might vary. We also considered an uncorrelated uptake of ACT rounds during MDA, meaning there is no relationship between the likelihood of receiving a future ACT round and having received a previous one. This can be an issue in areas where religious and/or cultural beliefs cause individuals to refuse any and all drug treatments, but if that is the case, then it would manifest itself at the time of the prevalence survey, when blood would have to be drawn. In practice, we concede that we may be unable to deploy MDA in whole villages due to these constraints, but in the absence of data we refrain from making any assumptions.

Here, we present a theoretical exploration of the potential impact of MDA strategies in different settings of the GMS, with special emphasis on the sensitivity of the predicted impact to logistical constraints, and transmission or population topologies. The ranges of parameters and distributions explored are meant to represent the current malaria situation in the GMS but need to be adjusted for application to specific areas/countries. In conclusion, we propose that mass drug interventions can be an invaluable tool towards malaria elimination in the right context. The model presented here predicts that an MDA’s success likelihood is bounded by the initial malaria prevalence and we elucidate how those chances can be improved through tailoring of implementation logistics. Although MDA is being revisited by the global community, very little attention has been paid to implementation logistics, and there seems to be no protocol adjustment across settings with completely different seasonality and human mobility patters, thus risking a sub-optimal MDA outcome.

## Methods

We developed an individual based, discrete time, spatially explicit, stochastic model, with mosquito population dynamics and human population movement. The flow diagram in Figure 1A describes the natural history of malaria infection in the human population. Details of how the dynamics of malaria transmission, human mobility and interventions are simulated are provided in the simulation protocol section below. All model parameters are presented in Table 2 along with their respective references when applicable. The simulated synthetic population mimics the demographics of a set of 1000 villages in SE Asia, with the distribution of villages over space, village sizes and age distribution of people likely not applicable to African settings. They should be generic enough to give a fair representation of rural settings in SE Asia. The parameter exploration presented here provides the limits of plausibility in terms of how people are expected to move, what levels resistance and malaria prevalence are at, and how spatially heterogeneous transmission is.

**Table 2.** List of parameters used in the model.

| # | Name | Description | Value | Reference |
| --- | --- | --- | --- | --- |
| 1 | $V$ | Set of villages | $\vert V\vert =1000$ | |
| 2 | $N$ | Set of individuals across all villages | $\vert N\vert =314,795$ | - |
| 3 | $ml$ | Proportion of males in the population | 0.48 | (43) |
| 4 | $maxage$ | Maximum age | 80 years | - |
| 5 | $beta$ | Biting rate | variable | - |
| 6 | $previ$ | Initial proportion of infectious people | variable | - |
| 7 | $resit$ | Initial proportion of artemisinin resistant infections | variable | - |
| 8 | $netuse$ | Proportion of individuals that own an insecticide treated bed net | 0.5 | - |
| 9 | $itneffect$ | proportional decrease of individual susceptibility/infectiousness related to ITN usage | 0.2 | - |
| 10 | *ovstay* | Mean number of nights spent somewhere when undertaking short-term population movement | 3 | - |
| 11 | *crit* | Critical distance below which overnight stays somewhere other than your home are made very unlikely | 4 km | - |
| 12 | *timecomp* | Mean time to complete ACT routine treatment | 4 days | Best guess |
| 13 | *fullcourse* | Proportion that receives treatment full course | 0.8 | (44) |
| 14 | *covab* | Proportion of symptomatic cases that receive antimalarials | 0.6 | (44) |
| 15 | *nomp* | Relative probability of receiving treatment in a non-malaria post village | 0.1 | Best guess |
| 16 | *asymtreat* | Relative probability of receiving treatment without clinical symptom | $10^{-4}$ | - |
| 17 | *tauab* | Daily probability of receiving ACT in a village under MDA | 1/1.5 | - |
| 18 | *gamma* | Mean liver stage duration | 5 days | (45,46) |
| 19 | *sigma* | Mean time to infectiousness after liver emergence | 15 days | (47) |
| 20 | *mellow* | Mean duration of symptoms | 3 days | (48) |
| 21 | *xa0* | Daily probability of going below the minimum effective artemisinin concentration | 1/7 | (49) |
| 22 | *xai* | Daily probability losing the DHA effect as part of ACT | 1/3 | (50,51) |
| 23 | *xab* | Daily probability of going below the minimum effective piperaquine concentration | 1/30 | (50,51) |
| 24 | *xpr* | Daily probability of going below the minimum effective primaquine concentration | 1/2 | (52) |
| 25 | *delta* | Mean duration of a malaria untreated infection | 160 days | (46,53,54) |
| 26 | *imm_min* | Minimum clinical immunity period | 40 days | Best guess |
| 27 | *alpha* | Average permanence in each immunity level | 60 days | - |
| 28 | *phic* | Relative infectiousness of symptomatic infections compared to sub-patent ones | 1 | - |
| 29 | *mdi* | Mosquito daily probability of dying while infectious | 1/7 | (55) |
| 30 | *mdn* | Mosquito daily probability of dying while infected but not yet infectious | 1/20 | (55) |
| 31 | *mgamma* | Mean extrinsic incubation period | 14 days | (56) |
| 32 | *amp* | Amplitude of mosquito density seasonal variation | 0.6 | Best guess |
| 33 | *process* | Days needed to administer a full ACT course in one village | 4 days | optimistic guess |
| 34 | *rounds* | Number of drug rounds in an MDA campaign | 3 | Standard practice |
| 35 | *btrounds* | Number of days between drug rounds in an MDA campaign | 32 | Standard practice |
| 36 | *vcefficacy* | Vector control efficacy | variable | - |
| 37 | *cBroda* | Daily probability of clearing blood stage drug sensitive parasites with circulating dha | 1/5 | (57,58) |
| 38 | *cBrada* | Daily probability of clearing blood stage artemisinin resistant parasites with dha | $G\_cBroda * 0.27$<br>(0.05) | (59) |
| 39 | *clroda* | Daily probability of clearing infectious stage drug sensitive parasites with circulating dha | 1/3 | (57,58) |
| 40 | *clrada* | Daily probability of clearing infectious stage artemisinin resistant parasites with dha | $0.27 * G_{clroda}$<br>(0.09) | (59) |
| 41 | *cBrodab* | Daily probability of clearing blood stage drug sensitive parasites with circulating dha- piperaquine | 1/3 | (57,58) |
| 42 | *cBradab* | Daily probability of clearing blood stage artemisinin resistant parasites with dha- piperaquine | $0.27 * G_{cBrodab} + (1.0 - 0.27) * G_{cBrodb}$<br>(0.33) | (59) |
| 43 | *clrodab* | Daily probability of clearing infectious stage drug sensitive parasites with circulating dha- piperaquine | 1/3 | (60) |
| 44 | *clradab* | Daily probability of clearing infectious stage artemisinin resistant parasites with dha- piperaquine | $0.27 * G_{clrodab} + (1.0 - 0.27) * G_{clrodb}$<br>(0.126) | - |
| 45 | *cBrodb* | Daily probability of clearing blood stage drug sensitive parasites with circulating piperaquine | 1/3 | (61) |
| 46 | *cBradb* | Daily probability of clearing blood stage artemisinin resistant parasites with piperaquine | 1/3 | (61) |
| 47 | *clrodb* | Daily probability of clearing infectious stage drug sensitive parasites with circulating piperaquine | 1/20 | (62) |
| 48 | *clradb* | Daily probability of clearing infectious stage artemisinin resistant parasites with piperaquine | 1/20 | (62) |
| 49 | *clrodp* | Daily probability of clearing infectious stage drug sensitive parasites with primaquine | 1/1.5 | (52,63) |
| 50 | *clradp* | Daily probability of clearing infectious stage artemisinin resistant parasites with primaquine | 1/1.5 | - |

Spatial demographics for both mosquitoes and humans are implemented at the village level, *i.e.*, humans can only move between villages and transmission within each village follows a pseudo-homogenous process where each mosquito is equally likely to bite a given individual. We assume villages are transmission units that encapsulate the village itself and the surrounding farms/forest areas. This is clearly a simplification of reality that whilst structurally convenient, since the village is our programmatic intervention unit, also happens to reflect mosquito daily flight patterns. Given that people tend to move freely within their village and that mosquitoes can cover the distance between village and proximal farms over the course of a single day, we can reasonably assume that all humans in one village can potentially be bitten by any one mosquito in that village.

Malaria transmission in the Greater Mekong Sub-region (GMS) has been reported to be spatially heterogeneous (38–40). However, there is very little evidence as to the relative abundance of the main vector species and no quantification of densities exists at a large enough scale. Given the lack of data to inform the discrepancies in mosquito densities across different villages, we chose to explore three mosquito density distributions (*Mn*, *Ml*, and *Mp*). Two of those distributions represent extreme scenarios: one in which all villages have approximately the same biting rate (Gaussian distribution); another where the vast majority has very low biting rates with a few hotspots (Pareto). The third distribution illustrates a scenario possibly more applicable to most areas in which some villages have a quite high transmission intensity, but where most have low mosquito abundance. These distributions were chosen arbitrarily and are parameterised as follows:

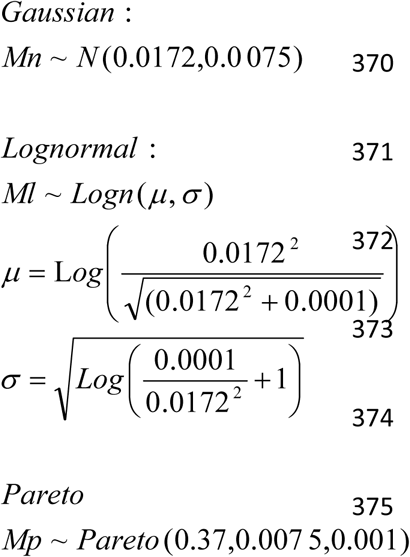

We take these distributions of mosquito density across villages as reference (*M* (*t* = 0)) and impose some seasonal variations to reflect the observed malaria incidence seasonal patterns. Mosquito density at time *t* in village *j*, for all mosquito density distributions is then given by:

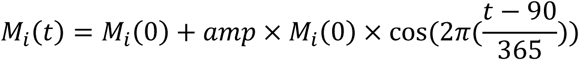

where *amp* reflects the amplitude of the seasonal fluctuation.

Transmission in a given simulation is characterised by a single extra parameter, the mosquito biting rate, which determines the vectorial capacity and is adjusted to reach a specific baseline malaria prevalence in the human population. The mosquito biting rate calibration was performed for each combination of transmission heterogeneity distribution, population mobility intensity and malaria mean prevalence. Thus, for each population mobility and mosquito density distribution, hundreds of model runs were performed until the desired mean malaria prevalence was reached with a specific mosquito biting rate within the first 5 years of simulation.

Population movement patterns and their importance for infectious disease transmission and emergence has recently garnered a lot of increased scientific interest, with new tools and analysis frameworks being developed for mobility inference (41, 42). Whilst census data and mobile phone data can help in proposing a connectivity network for a given region, the lack of general precision in questionnaires and the relative difficulty in capturing a lot of outside home overnight stays in mobile phone records, begs for a new source of data to resolve transmission relevant mobility patterns. Spatially explicit malaria models have in so far used gravity models to describe population movement, which is supported by some data (notably, day time travel data). Whilst we agree that a gravity model can be the most appropriate to describe some seasonal and long-term migration patterns, we argue that overnight stays a very short distance from your home are generally unlikely. It is more likely for someone to return home for the night if they are within a certain critical distance threshold (*crit*), instead of staying overnight somewhere else. We thus consider that daily population flow between villages is best characterised by a modified gravity model given by:

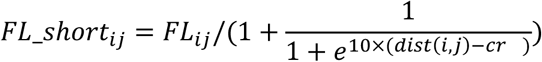

where *FL_ij_* refers to a standard gravity model presented below, and *dist* to the Euclidean distance between pairs of villages.

The frequency of general population’s short-term movements (very small number of nights spent somewhere other than their own village at a time) is given by the overall population mobility parameter – *mobility* – which is explored at length in the main text and assumes 2 extreme values (25 and 250 nights spent somewhere other than the home village, per year) and an intermediate one (125 days). The population is further partitioned into temporary (seasonal) or long-term migrant groups. Seasonal migration can only occur during a 3-month period. This roughly corresponds to the duration of crop seasons, at which time people typically go back to their village of origin to help their families harvest crops or for other economic/personal reasons. After 3 months they return to larger villages or surrounding cities following *FL_ij_*. Long term migrants only move between *a priori* defined (at random) economic hubs, comprising 2% of the target villages, spending an average of 6 months in each before moving to the next. These migration flows are characterised by a general gravity model:

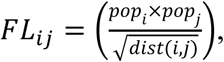

where *pop* refers to village population size.

We simulate malaria elimination strategies composed of MDA, a malaria village worker (VMW) network for improved case management, and an annual bed net distribution program. Villages are given an MDA of one full course (3 monthly rounds) of artemisinin combination therapy (ACT) plus one dose of primaquine, irrespective of their illness or infection status. Logistically, intervention teams sweep through all villages and give out ACTs without any prior screening, staying for a given number of days and then moving on to the next village. The details of how implementation logistics were incorporated into the simulation protocol can be found in the Simulation Protocol below.

When exploring the factors driving MDA outcome prediction, we explored all possible combinations of parameter values presented in Table 1, comprising 3888 sets of parameters. The model was run 100 times for each parameter set.

We also explored the layering of further intervention efforts on top of a global MDA initiative (with 3 ACT rounds), such as the implementation of indoor residual spraying (IRS) or vaccination. These control measures translate into a proportional decrease in infection risk for each individual living in a sprayed house or having received the vaccine. Thus, a transmission reduction efficacy of 0.10 means that the implemented vector control strategy reduces the number of infectious bites received per person per year by 10%. Whilst MDA is programmatically well defined, with a specific number of ACT rounds being deployed, it is less clear how vector control strategies are sustained over time. Thus, we ran simulations for a range of vector control efficacy/duration of intervention pairs and evaluated the proportion of simulations in which elimination is reached.

### Simulation Protocol

In this document, we give detailed description of the agent-based malaria simulation model results from which were discussed in the main text. We start by defining the interacting agents and their properties in the next section. Model processes and functions executed during the simulation are then documented in another two sections according to their positions in the simulation sequence.

#### Agents

The simulation model presented here is a multi-level agent-based model containing three interconnected groups of agents: Villages, Humans and Mosquitoes. Each agent has group-specific properties:

#### Villages

– Village ID
– Population size
– Location (Longitude, Latitude)
– Mosquito density
– Malaria post status / Date of establishment
– Current treatment strategy (whether the village is undergoing MDA)
– Number of administered ACT rounds Humans

#### Human ID

– Home Village ID
– Current Village ID
– Age
– Gender
– Susceptibility/infectiousness
– ITN usage (affects the susceptibility/infectiousness property above)
– List of infections (Each human entity can have several co-infecting parasites. Each parasite’s drug resistant status and maturity over time is tracked.)
– Transmission status
– Clinical status
– Immunity status (Immunity level and cumulative number of lifetime infections determined the probability of developing clinical symptoms upon infection)
– Treatment status
– Active circulating drugs (which drugs are circulating at effective concentrations in the person’s blood)

#### Mosquitoes

– Transmission status
– Infection carried (This property informs on the drug resistance of the parasites in the mosquito’s salivary glands)
– Current Village ID

### Model Initialisation

The developed malaria micro-simulation platform takes inputs from two CSV-formatted data files and a JSON-formatted configuration file. The two data files provide the model with a list of villages and a list of humans respectively. Each row of these two data files describes the properties (as listed in the previous section) of either a village or a human agent. Model-wide and process-specific, as opposed to agent-specific, parameters are given by the configuration file. Most parameters specified in Table 2 of the main text are associated with processes and functions (rather than individual agents), and therefore are given by the configuration file. In this section, we use the numbers in square brackets to refer to the associated parameter number whose description and value can be found in Table 2.

The initialisation process starts by processing the configuration file to get information such as the location of the other input files. Then a list of village agents and a list of human agents are created according to information given in the data files. Human agents are either randomly assigned a home village from all available villages or assigned to villages in the same order as they appear in the input data file. Once all agents have been created, the software initialises the parameterised-functions of the model with the information given in the rest of the configuration file.

### Human Properties

Note that whereas the configuration file and the village data file are mandatory, the data file for humans is optional to the model software. Given that the population of each village is known from the village data file, in the absence of a human data file, a set of human agents (*N* [2]) may be generated by the software according to each village’s population size. We now give details of the processes that are part of the initialisation process when properties of human agents are not specified by input.

#### Gender and Age

Age and gender information of human agents are generated according to parameterised distributions. The probability of a human agent being male is *ml* [3]. The age of a human is sampled from a discrete distribution specified by a vector of size *maxage* [4]. This vector is given by a csv file with *maxage* [4] integers.

#### Prevalence

The probability of a human agent being infectious is previ [6], and the probability of that infection being resistant to artemisinin is resit [7].

#### ITN Usage

Each human agent is assigned with an ITN with a probability of netuse [8]. Ownership of an ITN reduces the human agent’s susceptibility/infectiousness by itneffect [9].

#### Immunity

Each human agent is assigned two properties in relation to immunity, namely Cumulative Number of Exposures and Immunity Level. Both properties are set to 0 as default.

### Village Properties

#### Mobility Network

Geo-spatial human mobility amongst the population is a key element simulated in our model. Using the location and population information provided for each village, a complete graph (*FL*) is constructed linking all villages during initialisation. Let *M* denote the set of villages in the simulation, *FL* is constructed using a generic gravity model, with each edge denoting the flow of human movements between village *i,j* ∈ *V* as

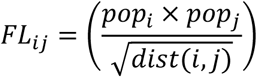

where *pop*_*i*_ denotes the population of village *i*, and *dist*(*i,j*) denotes the earth-surface distance between *i* and *j*.

### Mosquito Density

For each village, the mosquito density property allows the creation of the desired number of mosquitoes in each village. For simplicity sake, we chose not to create infectious mosquitoes during initialisation, but use a free parameter beta [5], the mosquito biting rate, to calibrate each simulation to the desired malaria prevalence.

### Implementation of malaria relevant dynamics

Once initialisation finishes, the main body of the model simulation starts. We assume time zero to be the 1st of January 5 years prior to the first malaria post establishment/MDA initiation.

The simulation protocols detailed in this section illustrate the flow of events and processes taking place each day. These dynamic processes are reduced to daily probabilities of occurrence, with events occurring if a randomly drawn, uniformly distributed number between 0 and 1, is lower than the likelihood of something happening on a given day. Thus, for each individual and for every simulated daily time step, a random uniformly distributed number is drawn for each possible transition process (e.g. whether the individual dies, is infected, is treated, etc), with that specific transition taking place if the number drawn is lower than the respective daily probability of occurrence.

### Human Population Dynamics

#### Birth and death

The probability for the death of a human agent is given by

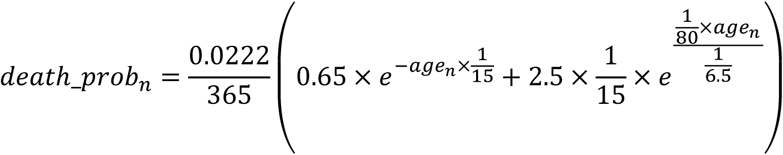

where *age*_*n*_ denotes the age property of the human agent *n* ∈ *N*. When a death event happens, a new agent is generated as a replacement. The new agent is placed in the same village where the death took place to keep population size a constant.

#### Short-term movement

Every human agent can display short-term mobility patterns, characterised by overnight stays in villages other than their home for a mean period of ovstay [10] days. For every agent who is currently located at their home village, the daily probability of such short-term movement is given by the factor Mobility in Table 1. Thus, the number of human agents embarking on short-term movement on a given day is

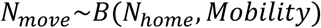

The destination of each agent’s movement varies and is determined using the mobility network *FL* constructed during initialisation. Let the flow of short-term movement between village *i,j* ∈ *V* be

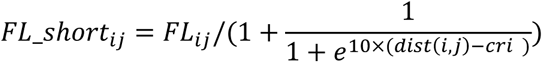

where *crit* [11] denotes the critical distance below which overnight stays at a village other than home are made very unlikely. The probability of a human agent to move from *home* to village *j* is

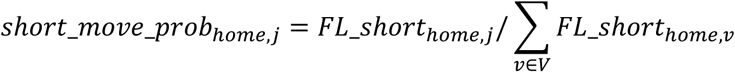

#### Clinical outcome

The likelihood of clinical symptoms brought on by a single infection is given by

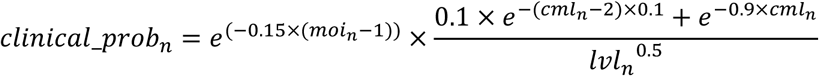

where *moi, cml* and *lvl* denote the multiplicity of infection, cumulative exposure to malaria and immunity level properties of the human agent respectively.

#### Treatment

The probability for a human agent to receive a full course of ACT treatment is dependent on symptomatology as well as the presence of a local malaria post. In a village with a malaria post, a human agent with clinical symptom would receive treatment with probability

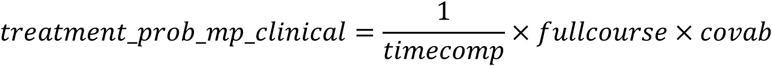

where *timecomp* [12], *fullcourse* [13] and *covab* [14] are described in Table 2. In a village with no malaria worker presence, this treatment probability is reduced by *nomp* [15]. Treatment rates of asymptomatic human agent is negligible as treatment is conditional on a positive RDT, which is very unlikely in sub-patent infections. The probability of a human agent with asymptomatic infection getting treatment is reduced by *asymtreat* [16].

In a village under MDA, the daily probability for a human agent to receive a round of ACT treatment is *tauab* [17].

#### Intrinsic incubation

Parasites emerge from the liver at a rate of 1/*gamma* [18], thus the liver stage takes on average *gamma* [18] days to complete

#### Gametocytaemia

Parasites start reproducing sexually, and thus generating gametocytes with 1/*sigma* [19] daily probability.

#### Clinical resolution

Malaria induced fevers gradually recede at a rate of 1/*mellow* [20], meaning that a person is feverish for *mellow* [25] days on average.

#### PK/PD

We describe waning drug efficacy over time through explicit daily probabilities of drug effect loss. Upon receiving treatment, each human agent will gradually lose the effect of both DHA and Piperaquine, according to *xai* [22] and *xab* [23] respectively. The single remaining drug will be lost at rates *xa0* [21] and *xab* [23] for Artesunate and Piperaquine respectively. Single dose Primaquine is lost at rate *xpr* [24].

Parasite killing rates depend on the person’s transmission status (*s*), with parasite clearance in not yet infectious people generally slower than that in individuals carrying gametocytes. Clearance of parasites with drug resistance phenotype *h* by drug *d* then follows

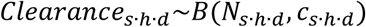

where *c*_*s.h.d*_ is an element of a 3-dimensional matrix of size |*S*| × |*H*| × |*D*|. *B* denotes a binomial distribution.

#### Recovery

Each infection in a human agent’s infection list has a daily probability of being naturally cleared given by 1/delta [25].

#### Immunity

One level of clinical immunity is gained by a human agent every time his infection list is emptied. Immunity loss starts imm_min [26] days after one level of immunity is gained. Immunity is lost at a rate of 1/alpha [27]. Therefore, each human agent is clinically immune an average of imm_min [26] + alpha [27] days. A loss in immunity prompts a reduction in immunity level and not the immune status per se.

#### Susceptibility/Infectiousness

Susceptibility was implemented as being solely dependent on ITN usage in these simulations. If someone sleeps under a bed net, their susceptibility to receive an infectious mosquito bite is reduced by itneffect [9]. Individuals sleeping under a bed net are also less infectious compared to people that don’t. Clinical status modulates infectiousness through phic [28], which determines the relative infectiousness of clinical malaria infections compared to sub-patent ones (here assumed to be 1).

#### Infection

Given the time dependent vectorial capacity of each village, we can extrapolate the number of mosquito bites landed on humans each day. We exclude all bites from non-infectious mosquitoes. For the remaining, the probability of causing a new infection in a human agent *n* is given by

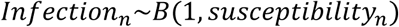

A resulting infection is then added to the human agent’s infection list and inherits the drug resistance phenotype of the infecting mosquito. The number of that agent’s cumulative number of exposures and multiplicity of infection is adjusted accordingly.

#### Mosquito Dynamics

Note that only infected and infectious mosquito agents exist in our model.

#### Survival

We assume adult female mosquito’s life expectancy to be mdi [29] + mdn [30] days on average. Meaning for each mosquito agent, its daily probability of dying while infectious is 1/mdi [29], and 1/mdn [30] while infected but not infectious.

#### Extrinsic incubation

Parasite development in mosquitoes takes an average of mgamma [31] days. Meaning for a mosquito agent, it takes mgamma [31] days from gametocyte infection to having sporozoites in the salivary glands and thus becoming infectious to humans.

#### Seasonality

Mosquito density at time *t* in village *i*, for all mosquito density distributions follows an annual seasonal cycle given by

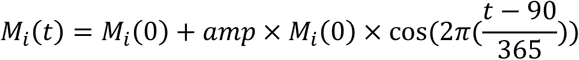

when only 1 seasonal peak is modelled in a year, and

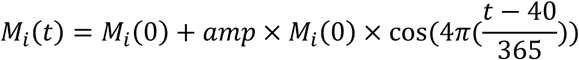

when 2 seasonal peaks are modelled in one year, with amp [32] denoting the amplitude of the seasonal variation of mosquito density.

#### Infection

Given the time dependent vectorial capacity for each village, we can easily extrapolate the number of mosquito bites landed on humans each day. For all bites handed out by mosquitoes that land on infectious humans we generate a new infection in the corresponding mosquito with the same resistant phenotype as a randomly sampled infection in the human’s infected list. There is no limit as to the number of infections a given mosquito can acquire during its lifetime. Their adult survivorship is quite limited though, making it uncommon for any female to reach multiparity.

### Interventions

#### Malaria Post

Malaria posts are established over time by placing a village malaria worker upon MDA initiation in each village.

#### MDA and Targeted MDA

In order to coordinate MDA teams to visit all villages without overlapping and repetition, a complete graph *VG* connecting all villages to each other is first constructed. The weight of the edge between village *i* and *j* is given by *dist*(*i*, *j*). This complete graph *VG* is then reduced to its minimum spanning tree form, denoted *MST*(*VG*), which we use to represent the road network connecting all villages.

Given that the MDA campaign includes *T* teams (Table 1), *T* starting locations are randomly selected from the nodes/villages of *MST*(*VG*). Then, *T* breadth-first-search algorithms are started from each of the *T* starting locations. These search algorithms run simultaneously in coordinated rounds. Each round, an algorithm is to add a village to its path. A village is added to the path of the algorithm which reaches it first. When an algorithm reaches a village that has been added/visited to the path of another algorithm, its searching continues until it finds an un-visited village in the same round. Once all villages have visited, all algorithms stop. Each algorithm’s path is used as the plan for each of the *T* MDA teams.

An MDA team stays in a village for process [33] days for each one of the rounds [34] number of ACT courses administered. There are btrounds [35] days between drug rounds.

A targeted MDA campaign is executed in a similar manner except that the initial graph *VG* is constructed only on the set of targeted villages.

#### Vector Control and Targeted Vector Control

Each village’s mosquito density property is reduced by 1-vcefficacy [36] during vector control. During a targeted vector control campaign, only a selected subset of villages’ mosquito density property is reduced.

## Acknowledgments

The authors would like to thank the members of the Malaria Elimination Task Force team and executive committee for their advice on the model structure and insights on the challenges of incorporating a large-scale targeted mass drug administration into a P. falciparum malaria elimination strategy.

---

**Figure 1 – figure supplement 1.**
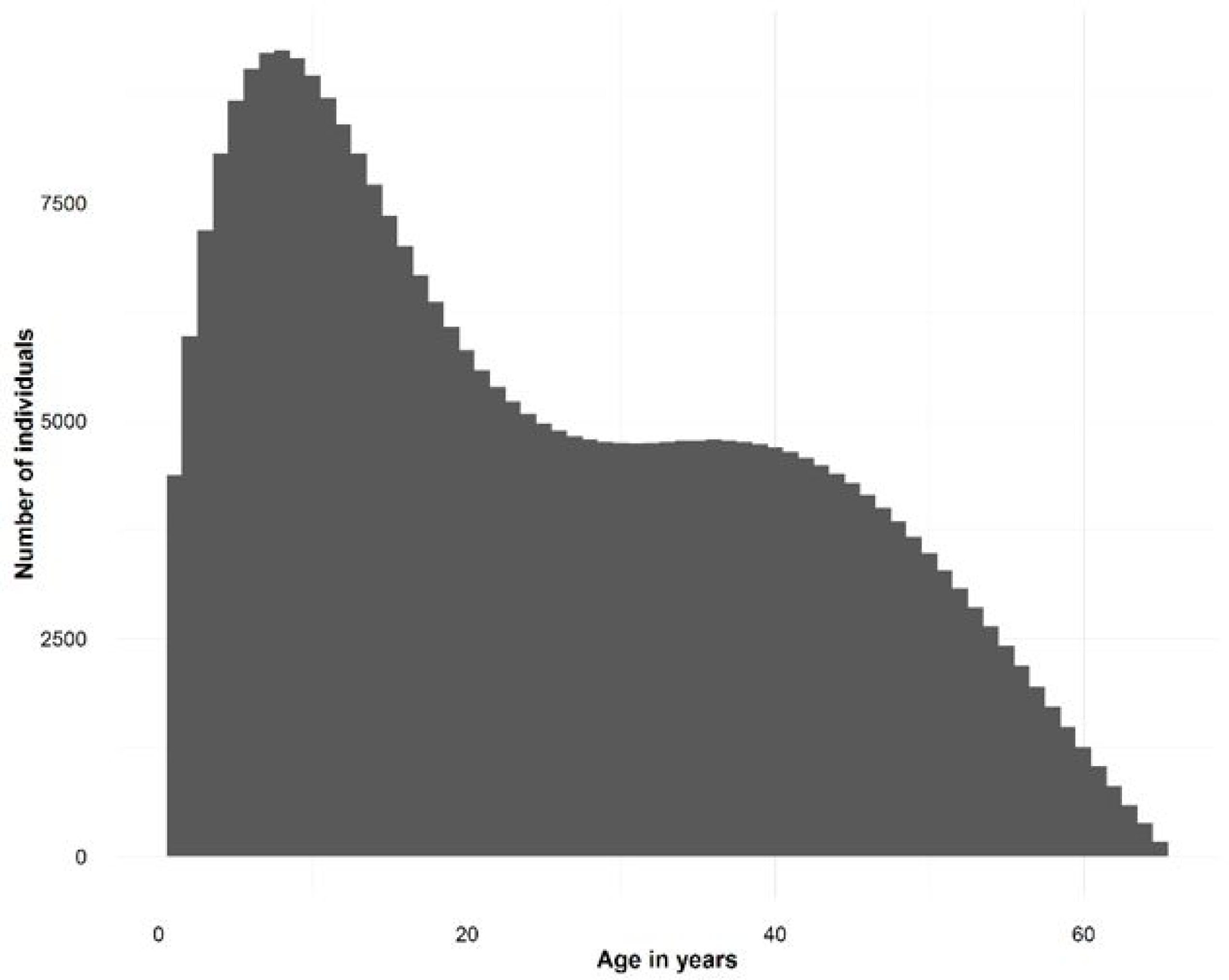
Synthetic population demographics. Depicts the age distribution of the simulated individuals.

**Figure 1 – figure supplement 2.**
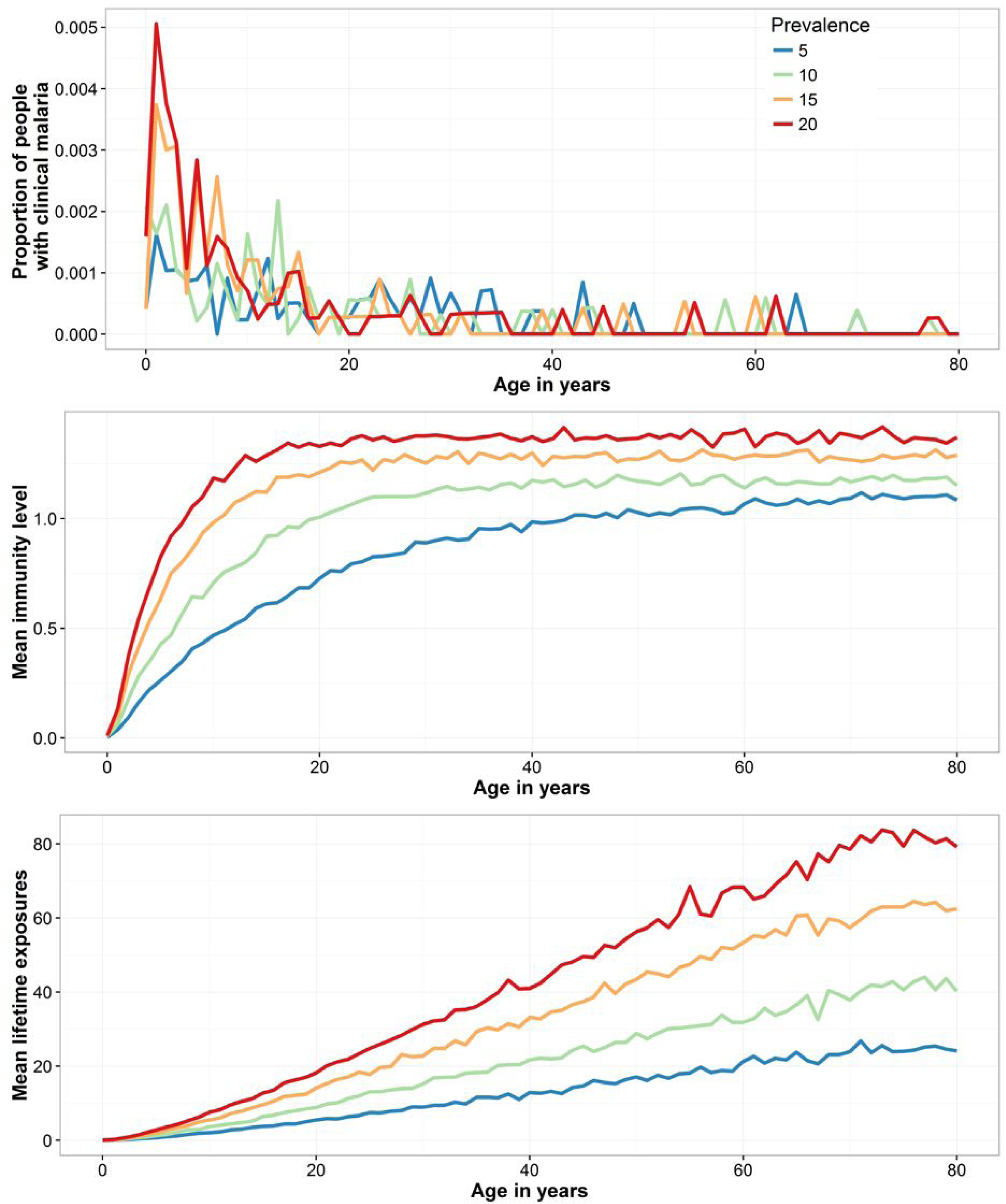
Clinical malaria age profiles and associated immunity and lifetime exposures. Displays epidemiological age profiles resulting from simulations employing the clinical immunity probability curves in Figure 1B. The several panels reflect how the function for clinical manifestations upon malaria infection is translated into clinical malaria age profiles for 4 different prevalence scenarios. The prevalence directly influences the number of lifetime exposures which interplays with clinical immunity loss to produce immunity level age dependent curves, which in turn regulate the resulting clinical age profiles.

**Figure 2 – figure supplement 1.**
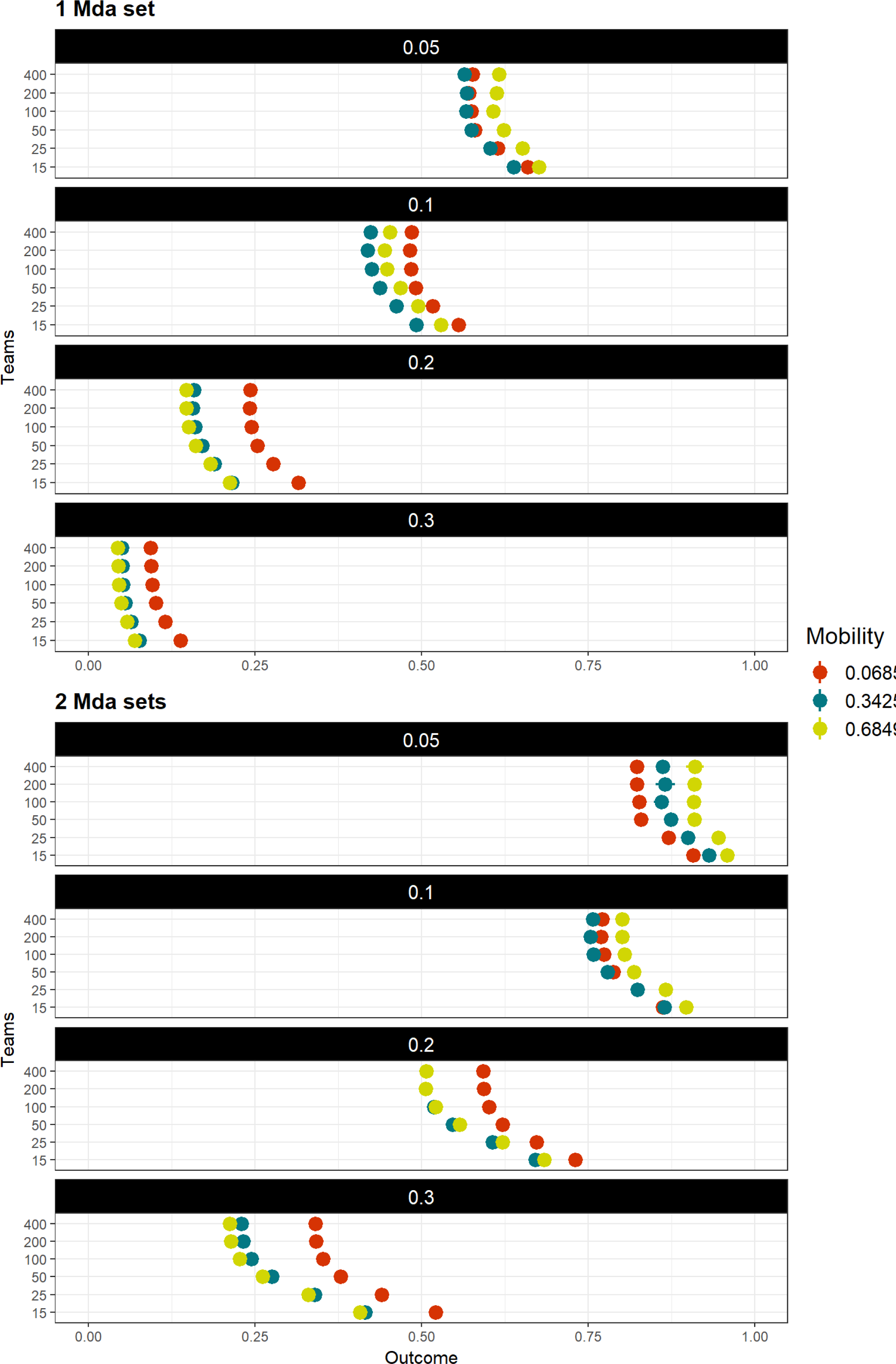
Multivariate model sensitivity analysis independent of transmission heterogeneity. Model outcome (measured as the proportional decrease in parasite prevalence) sensitivity to logistical (*Teams* – speed of coverage, and number of MDA campaigns), epidemiological (initial mean prevalence in the target area, given in black background over each panel), and behavioral (*mobility* – daily probability of spending nights in a place other than the habitual residence) parameters. Points and adjacent lines indicate the median and interquantile ranges (based on 100 simulations for each parameter set).

**Figure 2 – figure supplement 2.**
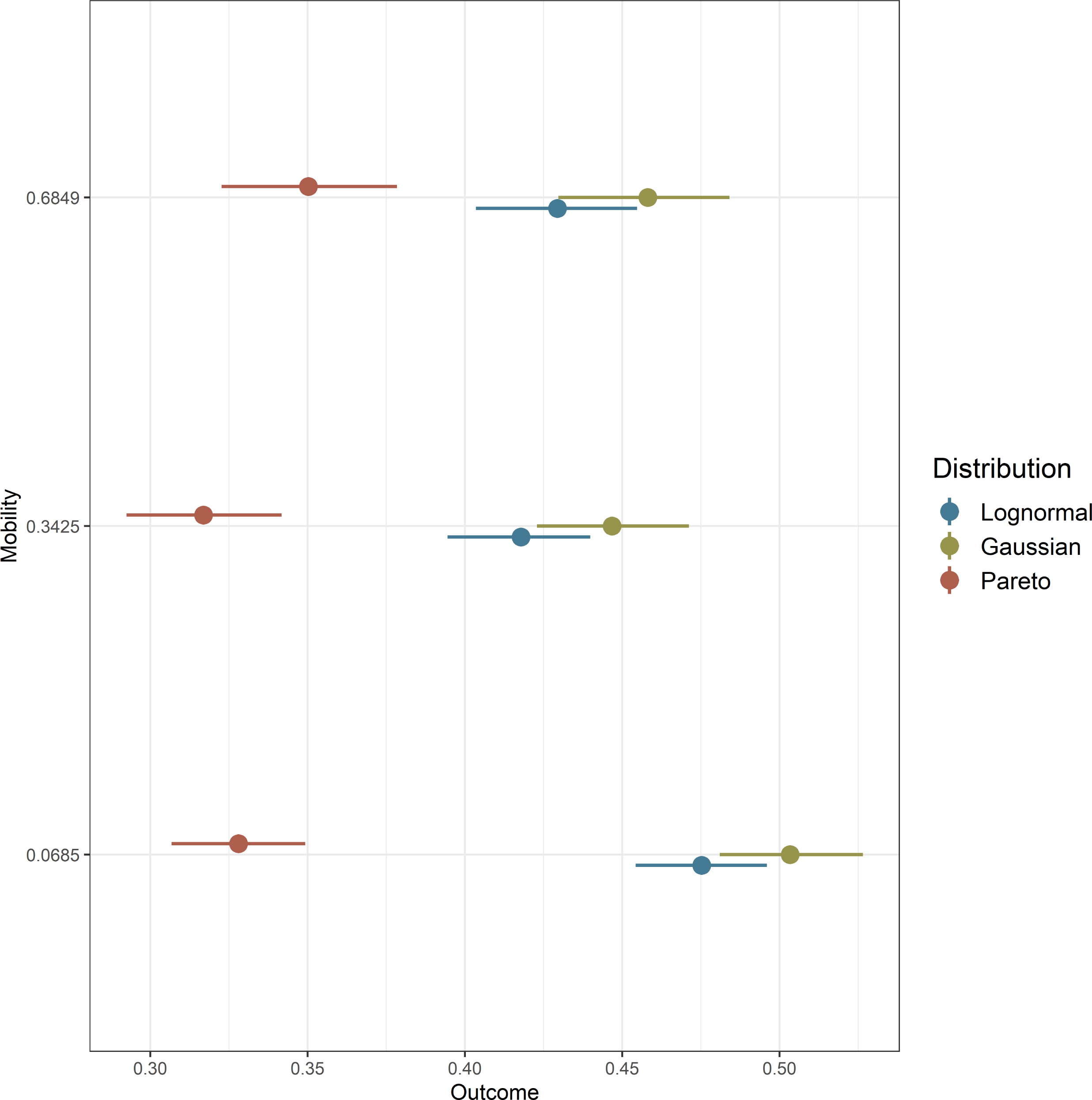
Interplay between population mobility and transmission heterogeneity. Intervention outcome sensitivity to spatial transmission heterogeneity in settings with different population mobility patterns. The vectorial capacity distributions associated with these simulations are displayed in Figure 1D. Points and adjacent lines indicate the median and interquartile ranges (based on 100 simulations for each parameter set).

**Figure 2 – figure supplement 3.**
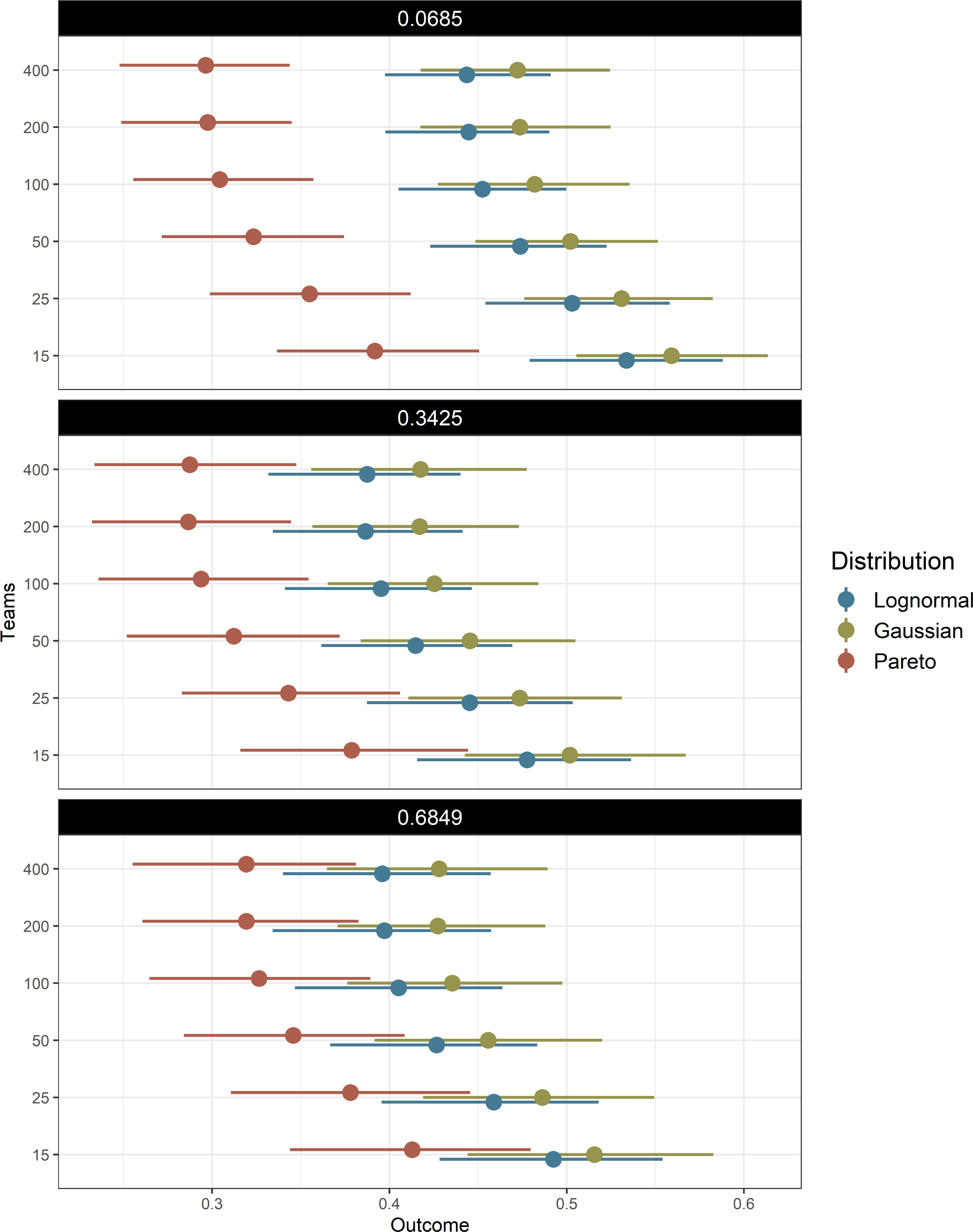
Interplay between logistics and human population topologies. Model predicted interaction of logistical MDA considerations (speed of coverage – teams) with different population mobility levels (given in black background over each panel) in determining intervention impact for the explored transmission heterogeneity distributions. Points and adjacent lines indicate the median and interquartile ranges (based on 100 simulations for each parameter set).

**Figure 2– figure supplement 4.**
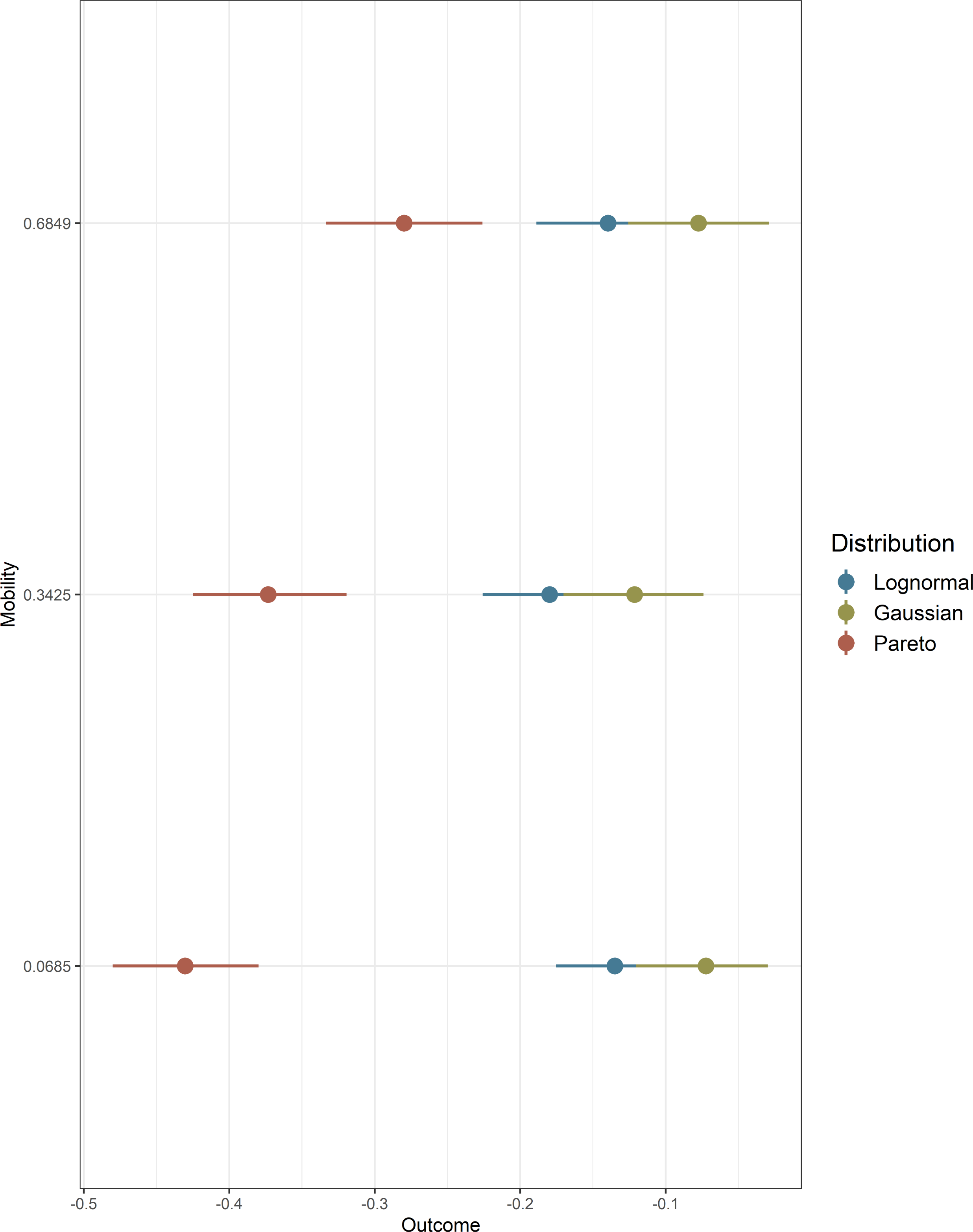
Factors influencing the spread of artemisinin resistance. In all similar to Figure 2, it shows how the different model parameters modulate the predicted proportional decrease in artemisinin resistance after MDA. Negative numbers then reflect a rise in artemisinin resistant infections from start of the campaign to the end of the simulation (a -100% value indicates a 2-fold increase in the number of resistant parasites).

**Figure 2 – figure supplement 5.** Population movement and mosquito distributions determine artemisinin resistance spread. Illustrates how human population mobility interacts with the transmission risk over space to modulate artemisinin resistance spread.

**Figure 3 – figure supplement 1.**
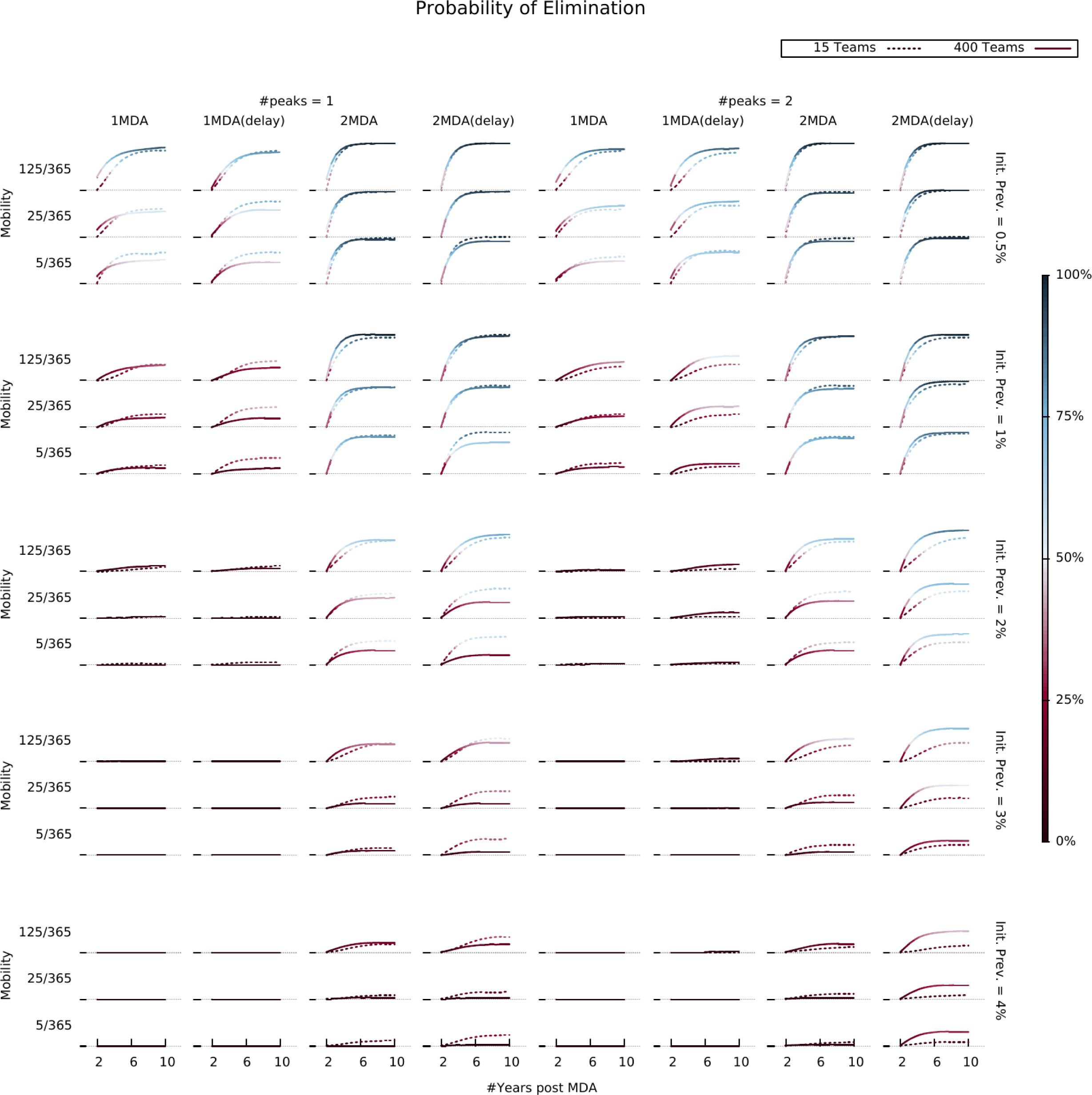
Elimination likelihood over time in different settings. Illustrates the interaction between human population mobility, seasonality patterns, and MDA operational parameters (number of teams and campaign start time). Colors indicate the proportion of simulations in which elimination was reached for each parameter combination, evaluated at different time points.

**Figure 4 – figure supplement 1.**
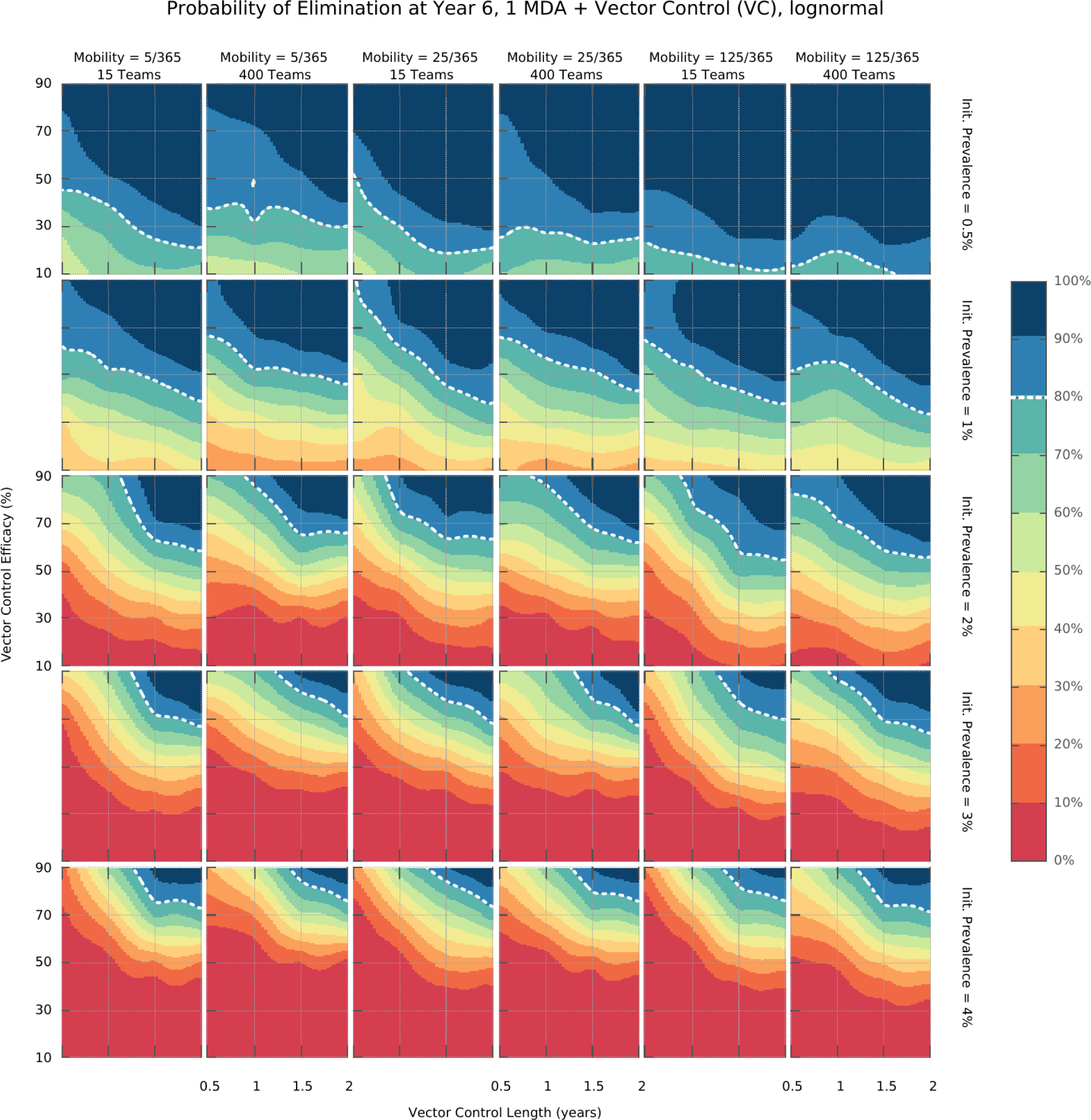
Integrated control elimination surfaces for the Log-normal distribution of transmission risk over space.

**Figure 4 – figure supplement 2.**
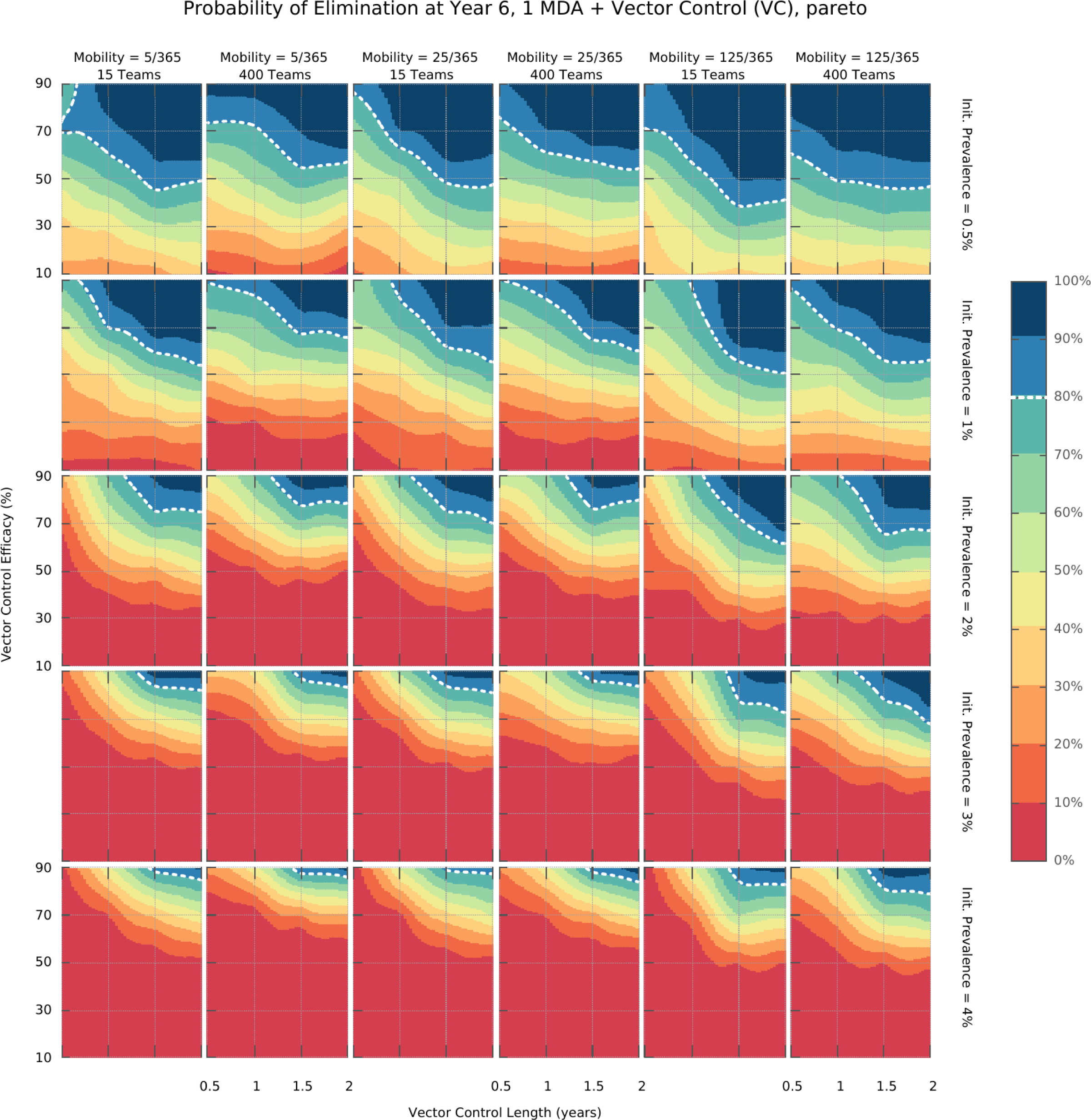
Integrated control elimination surfaces for the Pareto distribution of transmission risk over space.

**Figure 5 – figure supplement 1.**
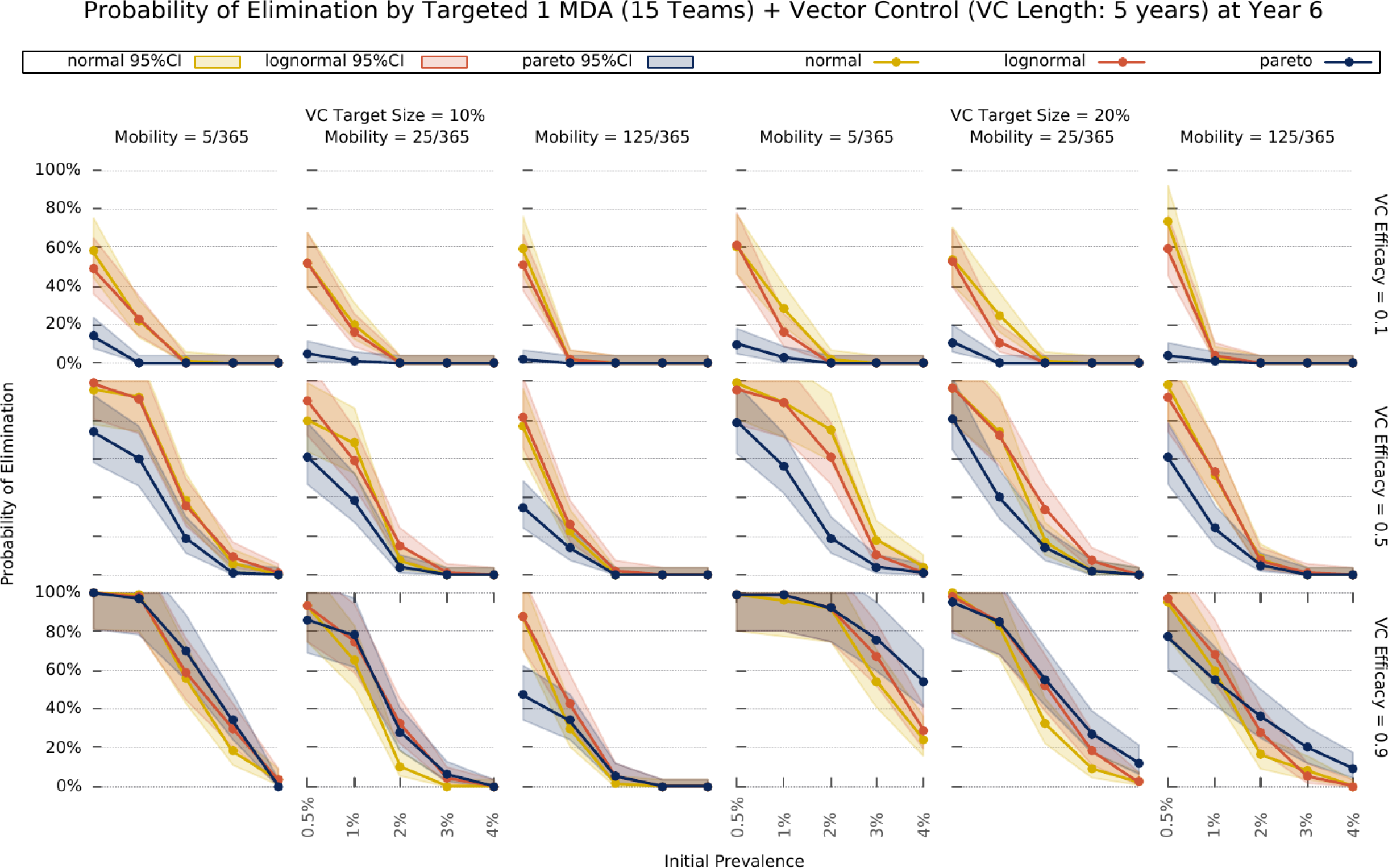
Elimination probability with a targeted approach. The same as in Figure 5, except that vector control is extended to last 5 years.

